# FMRP Regulates Neuronal RNA Granules Containing Stalled Ribosomes, Not Where Ribosomes Stall

**DOI:** 10.1101/2025.02.21.639553

**Authors:** Jewel T-Y Li, Mehdi Amiri, Senthilkumar Kailasam, Lily Drever, Jingyu Sun, Laura Bohorquez, Nahum Sonenberg, Joaquin Ortega, Wayne S. Sossin

## Abstract

Local protein synthesis is a crucial process that maintains local proteostasis in neurons. A large percentage of mRNAs translated in developing neurons are associated with stalled ribosomes. FMRP, the protein lost in Fragile X syndrome, is highly enriched in RNA granules that contain stalled ribosomes. Previous examination of ribosome protected fragments (RPFs) from stalled neuronal ribosomes identified sequences that match those found in mRNAs associated with FMRP, as identified by FMRP cross-linking immunoprecipitation (CLIP) (Anadolu et al, 2023, Journal of Neuroscience doi: 10.1523/JNEUROSCI.1002-22.2023). To investigate whether FMRP recognition of these sequences is important for determining where ribosomes stall on mRNAs, we examined RPFs isolated from P5 mice of both sexes that lack the FMRP protein. We found that the loss of FMRP had no significant effect on the proteins associated with neuronal stalled ribosomes, on ribosome structure, or the stalling sites (locations where RPFs accumulated). However, we observed a small, but significant decrease in the RPF levels from mRNAs previously shown to be associated with FMRP by CLIP in stalled ribosomes. Additionally, the number of neuronal RNA granules containing stalled ribosomes, as assayed by ribopuromycylation, decreased. Unlike neuronal RNA granules in WT neurons, the remaining neuronal RNA granules were resistant to reactivation. These results suggest a role of FMRP in neuronal RNA granules that contain stalled ribosomes, though loss of FMRP does not influence where ribosomes are stalled or the formation of stalled ribosomes.

## Introduction

Neurons are structurally unique cells with synapses–sites where neurons connect with each other– often located far from the cell body. For example, hippocampal pyramidal dendrites span an average of 13.5 mm (Ishizuka et al., 1995) while axonal tips can extend up to a meter away from the cell body (Debanne et al., 2011). To maintain and adapt the local proteome in response to local neuronal activity, neurons rely on the ability to produce proteins locally. Indeed, local protein synthesis has been shown to be critical for many aspects of neuronal function including growth cone guidance (Yoon et al., 2009), homeostasis (Holt et al., 2019), aspects of presynaptic firing (Wong et al., 2024) and certain forms of synaptic plasticity (Holt et al., 2019).

Local protein synthesis relies on coordinated mechanisms that couple mRNA transport with translational repression. Two major mechanisms have been proposed to regulate this process in neurons (Kiebler and Bassell, 2006; Sossin and DesGroseillers, 2006). In one mechanism, mRNAs are stalled during initiation and transported within RNA transport particles. In this case, the completion of initiation and onset of elongation depends on the removal of the repression mechanisms and availability of ribosomes, which must be transported separately. In contrast, mRNAs can be transported along with ribosomes when elongation stalls, with stalled ribosomes serving as the transport unit. Both mechanisms are likely to be utilized for neurons. While RNA transport particles appear to facilitate the transport of individual mRNAs (Batish et al., 2012), stalled ribosomes are found within large RNA granules that appear to contain multiple different mRNAs (Langille et al., 2019). The RNA binding protein (RBP) FMRP has been implicated in both types of transport (Richter et al., 2015).

Loss of FMRP causes Fragile X syndrome. In humans, the loss of FMRP results from expansion of a CGG repeat, leading to excessive methylation and transcriptional inhibition. Since CGG expansion is relatively common compared to de novo mutations and the gene is X-linked, Fragile X syndrome is a relatively common neurodevelopmental disorder and is the leading genetic cause of autism (Yu and Berry-Kravis, 2014). FMRP is an RBP and thus plays a role in various aspects of RNA biology. Several of these functions have been implicated in Fragile X syndrome, including FMRP’s regulation of miRNA repression, splicing, translation initiation, and translational elongation (Richter et al., 2015; Richter and Zhao, 2021). Another proposed cause of Fragile X syndrome is the loss of protein-protein interactions mediated by FMRP, including the direct regulation of ion channels (Deng and Klyachko, 2021).

A common finding in Fragile X models is that the loss of FMRP increases translation (Huber et al., 2002; Qin et al., 2005). This increase may result from the direct effects of FMRP on translation initiation or elongation, or from the loss of specific FMRP targets, which in turn lead to changes in signal transduction that ultimately increase protein synthesis (Bagni and Zukin, 2019). A major finding regarding how FMRP regulates translation is that, unlike most RBPs, FMRP exhibits strong association with the coding region in cross-linking immunoprecipitation (CLIP) studies, implicating FMRP in stalling elongation (Darnell et al., 2011). Consistent with a role in ribosomal stalling, loss of FMRP caused shifts in polysome profiles of CLIP-identified FMRP targets in Neuro2A translational extracts (Darnell et al., 2011). Additionally, elongation rates are also increased in mouse models of Fragile X, where FMRP is knocked out (Udagawa et al., 2013). While initial studies suggested that FMRP associates non-discriminatively throughout the coding region, more comprehensive bioinformatic analyses have revealed consensus sequences that were enriched in FMRP CLIP data (Ascano et al., 2012; Anderson et al., 2016). FMRP has also been shown to bind to specific motifs such as G quadruplexes and the kissing complex due to the specificity of the RNA-binding domains in the protein (Darnell et al., 2005), but the sequences enriched in FMRP CLIP data do not necessarily contain these motifs (Anderson et al., 2016).

Ribosomes from neuronal RNA granules have been enriched through sedimentation and biochemically and structurally characterized (Krichevsky and Kosik, 2001; Elvira et al., 2006; El Fatimy et al., 2016; Kipper et al., 2022; Anadolu et al., 2023). These ribosomes are stalled in the hybrid state (Kipper et al., 2022; Anadolu et al., 2023) and may possess other modifications that distinguish them from most ribosomes: For example, unlike most ribosomes, puromycylated peptides do not dissociate from neuronal stalled ribosomes (Anadolu et al., 2024). Additionally, anisomycin competes poorly with puromycin for puromycylation, a property not observed in most ribosomes (Anadolu et al., 2024). Examination of RPFs from these ribosomes have also shown differences from standard preparations. The RPFs are larger than expected and the motifs previously identified in FMRP CLIPs are enriched in the RPF sequences (Anadolu et al., 2023). It is not clear whether the enrichment of FMRP CLIP-associated sequences in RPFs from stalled ribosomes occurs because FMRP determines where ribosomes stall, or whether FMRP associates with stalled ribosomes that were already paused at these specific sequences.

To determine whether FMRP is directly involved in the formation of stalled ribosomes through sequence recognition, we compared RPFs from stalled ribosomes in mice lacking FMRP protein to those in WT mice of the same strain. We also tested whether the larger protected fragments could be reduced in size by higher nuclease treatment. Our findings show that stronger nuclease treatment does reduce the size of RPFs to a normal size. Despite its enrichment in RNA granules containing stalled ribosomes, FMRP does not appear to be important for the formation of stalled ribosomes since the loss of FMRP did not affect the association of other proteins with stalled ribosomes, the state of the stalled ribosomes, and, most importantly, where the ribosomes stall. However, using ribopuromycylation (RPM) to detect RNA granules containing stalled ribosomes (Graber et al., 2013), we found that the loss of FMRP decreased the number of RPM puncta. Thus, FMRP may contribute to stabilizing RNA granules.

## Results

### Comparing RNA Binding Protein Enrichment in WT and FMR1- RNA Granules

Stalled ribosomes are found in neuronal RNA Granules (Graber et al., 2013). Previous studies have demonstrated that in brain homogenates, a substantial number of ribosomes sediment in sucrose gradients used to separate monosomes from polysomes, and these have been suggested to be the ribosomes in neuronal RNA granules (Krichevsky and Kosik, 2001; Aschrafi et al., 2005; Elvira et al., 2006). This sedimentation was further optimized to separate the ribosomes found in the pellet from heavy polysomes (Fig. 1A) (El Fatimy et al., 2016; Anadolu et al., 2023). To determine whether the pellet from C57BL-6 mice is comparable to the pellet from rat brains in a previous study (Anadolu et al., 2023), we examined the protein distribution across all collected fractions from each species’ brain homogenate using Coomassie blue staining. The protein distribution was similar between rat brain and WT mouse brain with considerable amounts of proteins in both WT-mouse pellet and rat pellet (Fig 1B).

**Figure 1.**
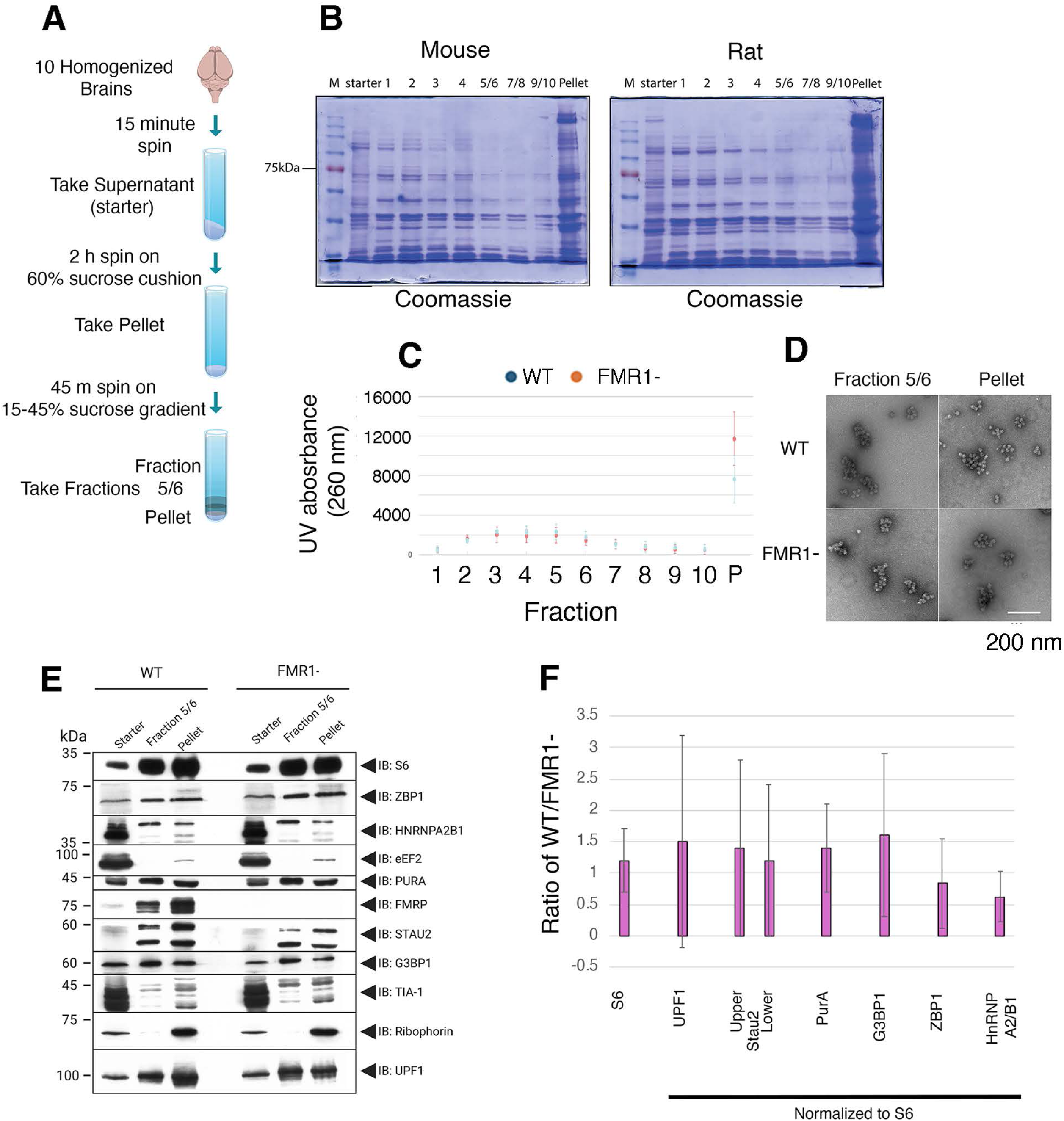
Characterization of Sucrose Gradient Sedimentation. A) A summary of the protocol for isolating the pellet and fraction 5/6 from mouse and rat whole brain homogenate using sucrose gradient fractionation. B) SDS-page stained with Coomassie brilliant blue showing the distribution of proteins from each fraction of the sucrose gradient from mouse and rat brains. Equal volumes of resuspended ethanol precipitates (fractions 1-10) and resuspended pellet were used. C) Average UV absorbance from fractions 1-10 and pellet from WT (Red) and FMR1- (blue) preparations (Error bars are S.D., N=3). D) Representative electron microscopy of WT and FMR1- ribosome clusters from fraction 5/6 and pellet. Scale bar is 200 nm). E) Representative immunoblots of fractions (as defined in A) for WT and FMR1- mice. For each blot, 1/100th of the Starter, 1/10^th^ of Fraction 5/6 and 1/20^th^ of the pellet fraction were loaded. F) Quantification of differences in enrichment between WT and FMR1-. Levels of proteins were determined from scans of immunoblots (see methods). Enrichment is defined by the ratio of level of protein (see Methods) between Fraction 5/6 and Pellet. For S6, the ratio of WT enrichment to FMR1- enrichment was calculated for each blot and the average calculated (N=8). For all the other RBPs, the WT and FMR1- enrichment were normalized to the S6 enrichment for that preparation and the normalized values were used to calculate the difference in enrichment between WT and FMR1- (UPF1, n=8, Stau2, n=8, PurA, n=3, G3BP1 (n=3), ZBP1 (N=3), hnRNPA2/B1 (N=3). Error bars are S.D. A one sample T-test against 1 was used to test significance with Bonferroni correction for multiple tests. No value reached p< 0.05.

We compared fractionation between WT and FMR1 knockout mice (FMR1-). FMR1-mice is the most common model for studying FMRP’s function (Richter and Zhao, 2021). There was no change in the distribution of UV absorbance in the fractions in the presence or absence of FMRP (Fig. 1C). We specifically compared fractions 5 and 6 from the gradient to the pellet to be consistent with previous studies (El Fatimy et al., 2016; Anadolu et al., 2023) comparing ribosomes in clusters that pellet and ones that do not. EM of fraction 5/6 and the pellet showed an abundance of ribosome clusters in both fractions in the presence or absence of FMRP (Fig. 1D). We examined the distribution of proteins in three fractions: the starting material after the first spin to remove non-soluble material, fraction 5/6, and the pellet, similar to the fractions examined in earlier studies (El Fatimy et al., 2016; Anadolu et al., 2023). We confirmed previous results from rat (Anadolu et al., 2023) that in WT mice FMRP (1.9 +/- 0.54 S.D, n=4, p<0.05 one sample T-test against 1) and UPF1 (1.5 +/- 0.57, S.D, n=8, p<0.05, one sample T-test against 1) were enriched in the pellet fraction vs fraction 5/6. As well, both Staufen 2 bands quantified (upper; 59 kDa and lower; 52 kDa) were enriched in the pellet fraction vs fraction 5/6 (59 kDa; 3.9 +/- 1.8, S.D, n=8, p<0.01, one sample t-test against 1; 52 kDa; 3.4 +/- 1.8, S.D, n=8, p<0.01, one sample t-test against 1).

We next compared the level of ribosomal protein S6 as a marker of ribosomes between WT and FMR1- mice. If FMRP was critical for the formation of the clustered ribosomes in the pellet, there should be fewer ribosomes in the pellet compared to fraction 5/6 in the absence of FMRP. However, we observed no significant difference in the ratio of S6 between these two fractions in the presence or absence of FMRP (Fig. 1E, quantified in Fig. 1F). We also compared the fractionation of proteins implicated in RNA transport in neurons between WT and FMR1-mice: UPF1 (Graber et al, 2017); Stau2 (Tang et al., 2001; Heraud-Farlow and Kiebler, 2014; Graber et al., 2017); Zip code binding protein 1 (ZBP1) (Huttelmaier et al., 2005), Pur alpha (Kanai et al., 2004), G3BP1 (Anadolu and Sossin, 2020) and hnRNP A2/B1(Gao et al., 2008; Lo et al., 2025). No significant differences in fractionation for any of these proteins were observed in the presence or absence of FMRP (Fig. 1E, quantified in Fig. 1F). TIA-1, a protein present in stress granules, but not neuronal RNA granules (Anadolu and Sossin, 2020; Sun et al., 2025) was mainly in the starter fraction. We examined eEF2 and confirmed previous results (Anadolu et al., 2023) that there was little eEF2 in either fraction 5/6 or the pellet under these conditions (Fig. 1E). We also examined the rough ER marker ribophorin. While previous results showed that secretory mRNAs showed low abundance in the pellet compared to most mRNAs, the secretory mRNAs were enriched in the pellet compared to fraction 5/6 (Anadolu et al., 2023). To determine if this is due to increased presence of rough ER in the pellet fraction vs fraction 5/6, we examined distribution of the rough ER marker ribophorin. We observe ribophorin protein in the pellet, but not in fraction 5/6 and this was not affected by FMRP (Fig. 1E).

### Loss of FMRP Does not Alter Anisomycin Puromycin Competition

Previous cryo-EM studies of ribosomes isolated from dense ribosome clusters showed that most of these ribosomes are in the hybrid A/P and P/E configuration (Kipper et al., 2022; Anadolu et al., 2023). Puromycin and anisomycin are both translational inhibitors that bind to the A site of the ribosomes (Hansen et al., 2003; Garreau de Loubresse et al., 2014). Due to their overlapping binding sites, anisomycin inhibits puromycin from binding to the ribosomes when both are present. However, we previously showed that puromycylation of neuronal ribosomes in cell culture and puromycylation of ribosomes in the pellet from rats are resistant to anisomycin inhibition (Anadolu et al., 2024). This suggested that the binding site for these translational inhibitors is altered in the neuronal stalled ribosome, either due to the enrichment of the hybrid state or due to other differences in these ribosomes.

We performed these experiments on the ribosomes that were sedimented through the sucrose gradient (Fig 2A). We first validated that puromycylation in non-neuronal ribosomes, in this case- rat liver ribosomes, were not resistant to anisomycin. Indeed, we saw a total inhibition of puromycylation by anisomycin in sedimented rat liver ribosomes (Fig 2B) much like the effect previously seen on HEK cell culture (Anadolu et al., 2024). This indicated that neither homogenization in our buffer nor high centrifugation is sufficient to confer puromycin’s resistance to anisomycin and further confirmed the neuronal specificity of this ribosomal state. If FMRP was important for generating the state of the ribosome required for anisomycin-resistant puromycylation, we should observe less resistance to anisomycin in FMR1- mice. We found that resistance of puromycylation to anisomycin was not different in ribosomes from the pellet of WT and FMR1- mice (Fig. 2B, quantified in Fig. 2C). Thus, FMRP does not seem to be important for the formation of this state of the ribosome.

**Figure 2.**
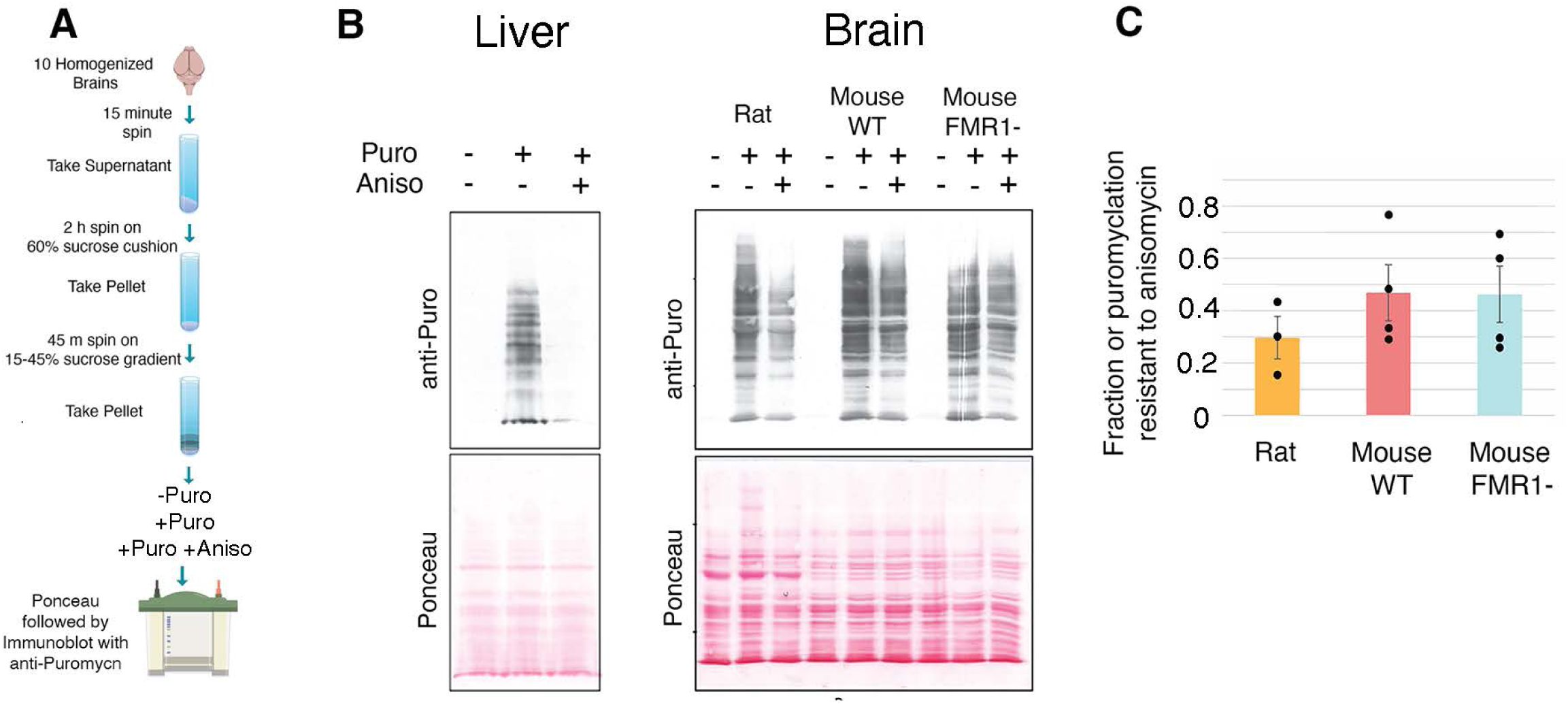
Loss of FMR1 does not affect anisomycin-puromycin competition. A) A summary of protocol for isolating the pellet and treating with puromycin. B) Representative immunoblot stained with antibodies to puromycin (anti-puro) (Top) to showcase the inhibition of puromycylation (100 uM) by anisomycin (100 uM) in liver polyribosomes (left) or comparing the pellet from Rat, Mouse WT and Mouse FMR1- (Right). Bottom: Corresponding membrane stained with Ponceau before immunoblotting. The liver experiment was replicated twice with similar results. C) Quantification of the amount of puromycylation resistant to anisomycin (puro +aniso/ puro) in Rat (N = 3), WT mouse (N = 4), FMR1- (N = 4). All groups are insignificant from each other (one way ANOVA, p > 0.05).

### Effects of Digestion on RPF Size

To explore if the stalling sites of the ribosomes are altered by the loss of FMRP, we examined RPFs (Fig. 3A). RPFs generated from the rat pellet were generally above 35nt (Anadolu et al., 2023), while RPFs have a canonical size between 28nt and 32nt (Ingolia, 2014). It was unclear whether the extension at the 3’ end from the RPF generated from the pelleted rat ribosomes was due to an extended conformation of the ribosome to protect a larger fragment or due to altered nuclease resistance in this region (Anadolu et al., 2023). Thus, we investigated whether this extended region could be cleaved off with more effective nuclease treatment. To determine whether the extending nucleotides could by removed by increasing nuclease efficiency, we digested at room temperature rather than 4°C, and in addition, the concentration of RNAse was adjusted to the sample RNA concentration (see Methods).

**Figure 3.**
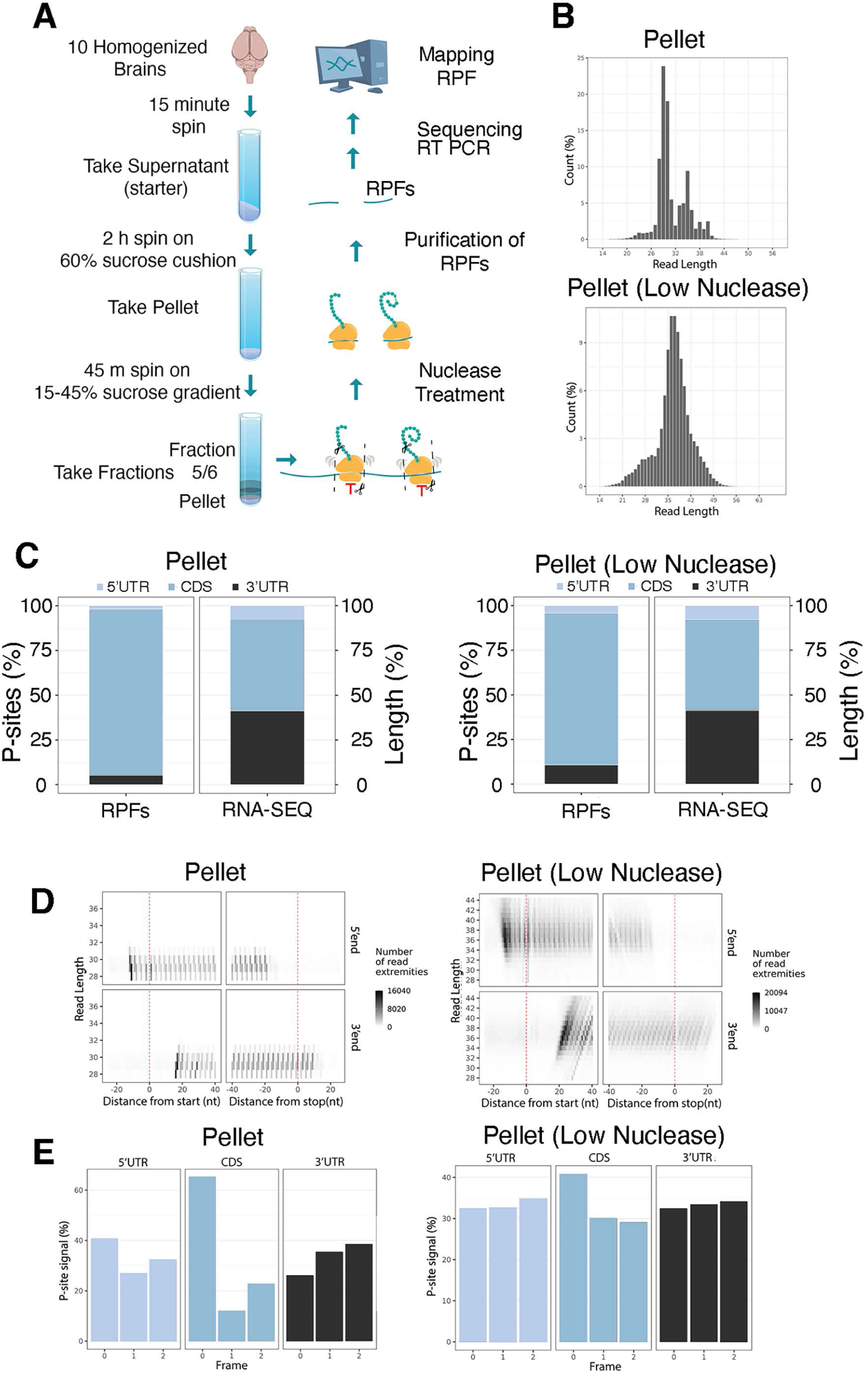
Higher Nuclease Reduces Size of RPFs in WT Pellet. A) A summary of the protocol for the RPF procedure. B) Size distribution of normalized footprint reads from the WT Pellet fraction under standard or low nuclease treatment. C) Representative image for read coverage for WT Pellet fraction either with standard or low nuclease treatment) UTR, untranslated region; CDS, coding sequence for both RPFs and RNA-seq libraries. D) Representative image for the number of read extremities (shading) for each read length (Y-axis) based on the distance from start(left) to stop(right) with the 5’ end (top) and 3’ end (bottom) for the WT pellet fraction with either standard or low nuclease treatment. E) Representative image for the periodicity statistics for RPFs in different regions of the mRNA for standard or low nuclease treatment. Although the representative images above included only one replicate, similar results were observed across all three replicates.

We replicated the increased size of RPFs in mice using the low nuclease protocol. However, RPFs generated by the new nuclease treatment decreased the RPFs to around 29-30 nt (Fig 3B). While there was still a portion of RPFs that stayed around 35 nt (Fig 3B), the majority of these RPFs are from noncoding mRNAs (Extended Data Table S3-1). The new nuclease treatment also showed a higher percentage of RPFs in the coding region (CDS) (Fig. 3C), no extension on the 3’ end (Fig. 3D) and a higher periodicity (Fig. 3E). These results were consistent across all libraries (Fig. S3-1, S3-2, S3-3). For all analysis below, we use only the RPFs from the new (standard) nuclease treatment.

### Higher MgCl_2_ Concentrations did not Affect Ribosomal Structure

An additional change from the previous protocol was in the concentration of MgCl_2._ Most ribosome isolation protocols use 10 mM MgCl_2_, but the previous protocol had a more physiological concentration of 2.5 mM MgCl_2_. Higher levels of MgCl_2_ in most protocols that purify ribosomes are presumably due to the finding that bacterial ribosomes dissociate in lower concentrations of MgCl_2_, but this has not been observed in eukaryotic ribosomes. A cryo-electron microscopy (cryo-EM) characterization of RNase I-treated pellet fraction under this low magnesium concentration (Anadolu et al., 2023) found that 85% of the ribosomes contained in the pellet exhibited tRNA molecules in hybrid A/P and P/E state. The remaining 15% of the ribosome population in this sample contained a tRNA in the P/P state. To test whether increasing the magnesium concentration had any effect on the percentage of ribosomes in hybrid A/P and P/E state, we repeated the cryo-EM and image classification analysis (Fig. S4-1) after the purification was performed in a buffer containing 20mM Tris-HCl, pH 7.4; 150 mM NaCl; 10 mM MgCl_2._ We found that high MgCl_2_ did not impact the percentages of hybrid A/P and P/E state (83%) and P/P (17%) state ribosomes (Fig. 4A). The cryo-EM maps for the hybrid and P/P state ribosomes were refined to 2.7 Å and 3.2 Å, respectively (Fig. S4-2 and Table S4-1).

**Figure 4.**
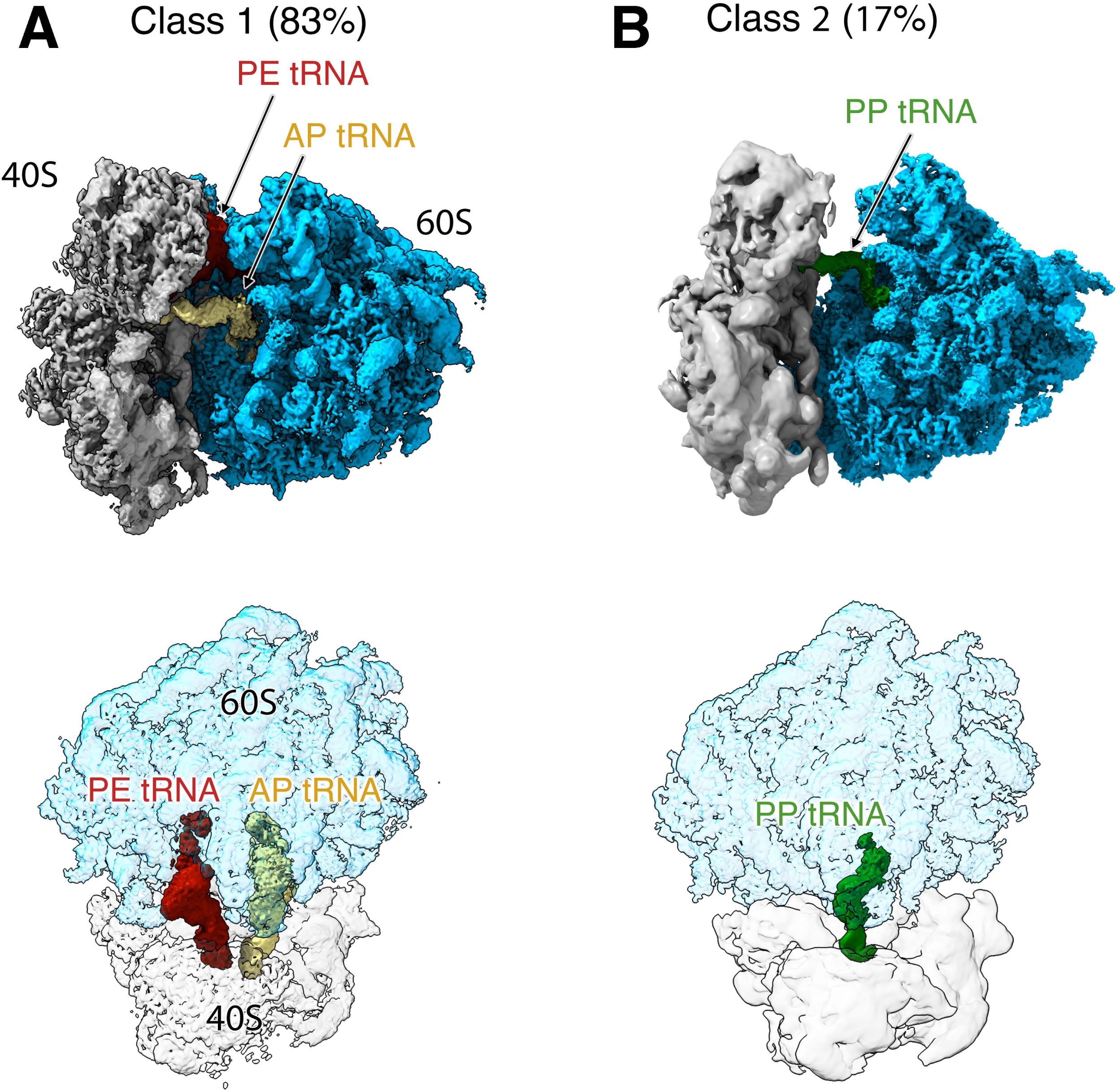
High Magnesium Buffer does not Affect Ribosome structure. Composite cryo-EM maps of class 1 (A) and class 2 (B) 80S ribosomes found in the pellet after purification in high magnesium buffer and RNase I treatment. The top panels show a side view of the two classes of ribosome particles contained in the sample. The bottom panels show top views of the same cryo-EM maps. The 40S and 60S subunits are shown as transparent densities to facilitate visualization of the positions of the tRNA molecules in each class.

### Characterization of the RPFs of mRNAs in WT and FMR1- Pellet through DEG, GO and GSEA Analysis

To analyze the mRNAs in the WT and FMR1- pellet, we mapped the data from RNA-seq and the RPFs to their corresponding mRNAs (Fig. 5A), similar to our previous analysis of RPFs from rat (Anadolu et al., 2023). PCA analysis shows that the variation between samples was similar to the difference between WT and FMR1- and that there was a large difference between the sequences in the RPFs and RNA-seq (Fig. S5-1). To compare the samples across different biological preparations, we calculated the reads per kilobase per million mapped reads (RPKM), which normalized the mRNA count against the total mapped count and gene length (Extended Data Table 5-1). We use the term abundance for the RPKM of RPFs found in the pellet fraction. The number of RPFs normalized against total mRNA obtained from whole brain homogenate with RNA-seq, often termed translational efficiency, was also calculated as the log(fold change) log (FC) (Extended Data Table 5-2). We refer to this measure as ribosome occupancy (Anadolu et al., 2023) since the subject of this study is stalled ribosomes and thus the number of ribosomes/mRNA is related not only to translational efficiency, but also to stalling. Lastly, we normalized the RPKM of the RPFs in the pellet against the RPKM of RPFs in fraction 5/6 as log (FC) to determine the extent to which mRNAs are selectively found in the pellet fraction (Extended Data Table 5-2). We call this value enrichment (Anadolu et al., 2023) to reflect enrichment in the pellet fraction.

**Figure 5:**
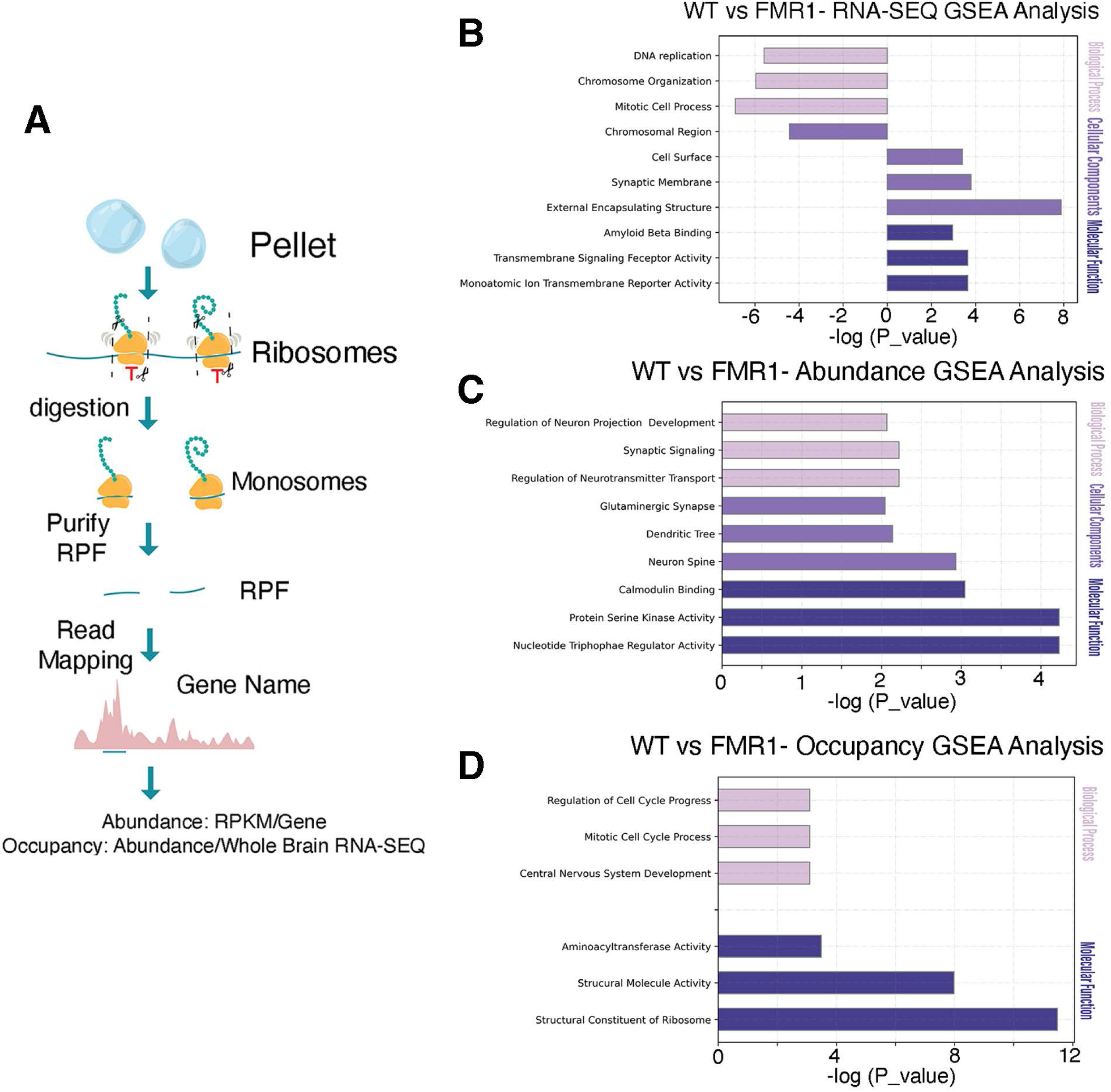
GSEA analysis WT and FMR1- RNA-seq and RPFs from pellet fraction. A) Description of RPF isolation and mapping. Results of Selected GSEA sets significantly affected by the loss of FMRP from analysis of B) RNA-seq, C) Abundance and D) Occupancy. Increases in FMR1- vs WT are to the left and decreases to the right.

We used DEG analysis to determine differences in RPFs beetween WT and FMR1-. Only a few differences were observed between RKPM values in WT and FMR1- RNA-seq samples (Extended Data Table 5-2). Similarly, there were very few differences using DEG analysis for abundance, occupancy or enrichment between WT and FMR1- (Extended Data Table 5-2). GO analysis of the top 500 abundant or occupied mRNAs gave similar results in WT and FMR1-mice and these results (increased abundance of cytoskeletal mRNAs and increased occupancy of mRNAs encoding proteins involved in RNA processing) were similar to previous results in rats (Figure S5-2; Extended Data Table 5-3) (Anadolu et al., 2023).

As another way to compare mRNAs between the two groups we used gene set enrichment analysis (GSEA) comparing WT and FMR1- samples (Fig. 5A, Extended Data Table 5-4). In the RNA-seq analysis, this revealed several gene sets whose levels increased or decreased upon loss of FMRP (Fig. 5B, Extended DataTable 5-4). This analysis revealed a decrease of the abundance of gene sets in FMR1- samples (Fig. 5C). GSEA sets identified as decreased in abundance in the pellet included postsynapse, neuron, spine, dendritic tree and protein kinases (Fig. 5C, Extended Data Table 5-4). There were also some gene sets that had decreased occupancy including cell component structural component of the ribosome, and nervous system development. There were no GSEA sets identified as changed in enrichment (Extended Data Table 5-4). We determined whether the modified mRNAs identified in the GSEA analysis were enriched for FMRP targets as defined by RNA CLIP studies (Darnell et al, 2011). Approximately 4% of the total mRNAs are FMRP targets and approximately this percentage of the gene sets identified in RNA-seq or measuring occupancy matched this expected number (Extended Data Table 5-4). In contrast, the gene sets decreased in abundance by the loss of FMRP had about 30% FMRP target mRNAs in the mRNAs changed. (Extended Data Table 5-4). This suggested that there may be a decrease in the levels of FMRP target mRNAs in the pellet.

### Comparison of the mRNAs in the WT and FMR1- to Selected Datasets

Previously, we showed that specific mRNA subsets are enriched in the pellet (Anadolu et al., 2023). We first examined the the abundant mRNAs associated with ribosomes resistant to ribosome runoff (Shah et al., 2020) (Extended Data Table 6-1). Similar to previous experiments with rats (Anadolu et al., 2023), the mRNAs resistant to run off were significantly abundant, occupied and enriched in both WT and FMR1- pellet RPFs (Fig 6). Next, we investigated FMRP associated mRNAs (FMRP targets) through assessing mRNAs identified by cross-linking FMRP with mRNA (Darnell et al., 2011; Maurin et al., 2018) (Extended Data Table 6-1). We found that FMRP targets were highly abundant and enriched in the WT and FMR1-pellet (Fig 6A, Fig 6C, Fig 6E), but the increase in occupancy was less than that observed for the mRNAs resistant to run-off. The increase in occupancy compared to total mRNAs was even less in the FMR1-samples.

**Figure 6:**
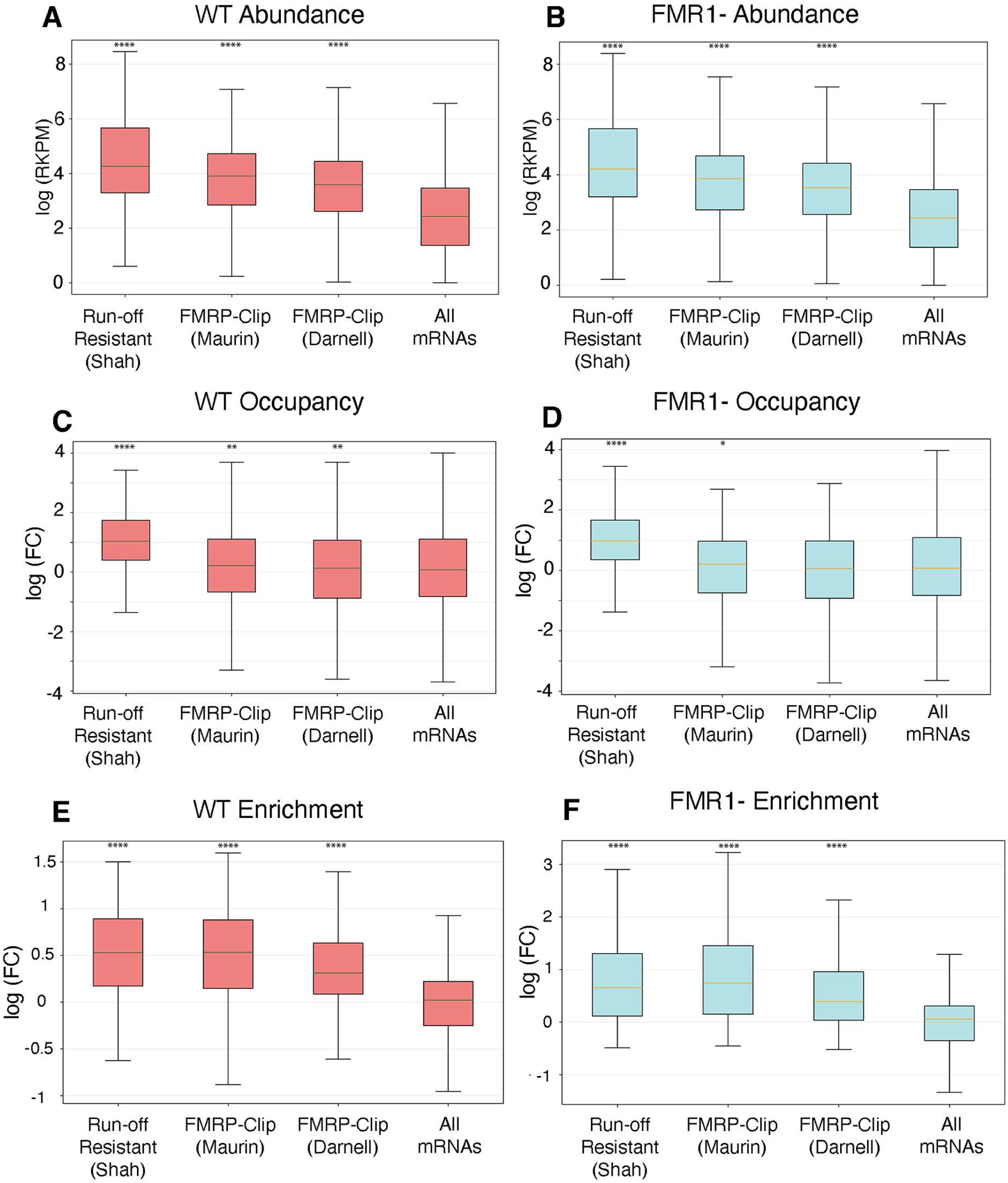
Comparison of Putative Stalled mRNAs vs Total mRNAs in RPFs from the Pellet of WT and FMR1- mice. Comparison of mRNAs associated with ribosome resistant to initiation inhibitor mediated run-off (Shah et al., 2020) and FMRP-CLIPped mRNAs (Maurin et al., 2018; Darnel et al., 2011) to all other mRNAs. A) WT Pellet abundance, B) FMR1- Pellet abundance, C) WT Pellet Occupancy, D) FMR1-Pellet Occupancy, E) WT Enrichment, and F) FMR1- Enrichment. As the N of the All mRNAs (13079) were much larger than for the selected groups: Shah (185), Maurin (264), Darnell (757) (Extended Data Table S6-1) a random set from the total RNA group that matched the number in the selected group was generated for a two-tailed Welch’s T-test with Bonferroni correction for multiple T-tests. The median P value from 10 random sets was used (Extended Data Table S6-1). (****, p<0.0001, ***, p<0.001, **, p<0.01, *, p<0.05).

To better understand this change after the loss of FMRP, we next examined the specific effect of the loss of FMRP on these subsets of mRNA by calculating the fold change in abundance, occupancy, and enrichment of these specific mRNAs between WT and FMR1-pellets compared to changes in all mRNAs (Fig. 7; (Extended Data Table 7-1). This revealed a significant decrease in the abundance, occupancy and enrichment of the FMRP targets. Strikingly, while there was also a decrease in the abundance and enrichment of mRNAs resistant to ribosome runoff, their occupancy did not change after the loss of FMRP. It should be noted that the decreases of FMRP targets in the FMR1- pellet were small (Fig. 7) and only significant when looked at in the whole subset. Individual mRNAs in this group were not found to be significantly different in the DEG analysis (Extended Data Table 5-2).

**Figure 7.**
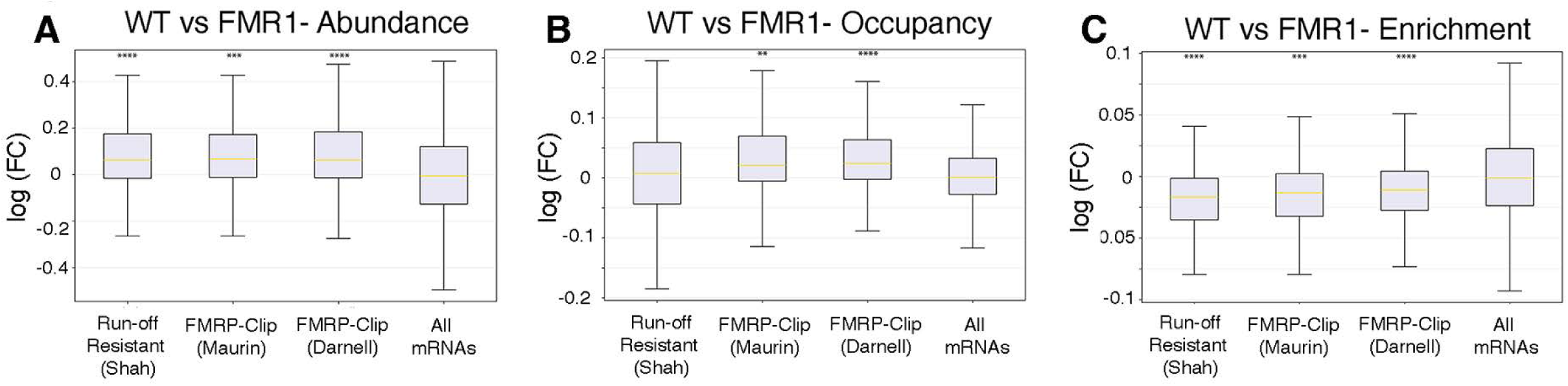
Comparison of Putative Stalled mRNAs between WT and FMR1- mice. Comparison of mRNAs associated with ribosome resistance of initiation inhibitor run-off (Shah et al., 2020) and FMRP-CLIPped mRNAs (Maurin et al., 2018; Darnel et al., 2011) to all other mRNAs. The fold change between WT and FMR1-was calculated using DEG (see Methods). These fold changes were then compared between the selected groups and All mRNas for A) Pellet abundance, B) Pellet occupancy, C) Pellet enrichment. As the N of the All mRNAs (13079) were much larger than for the selected groups: Shah (185), Maurin (264), Darnell (757) (Extended Data Table S7-1) a random set from the total RNA group that matched the number in the selected group was generated for a two-tailed Welch’s T-test with Bonferroni correction for multiple T-tests. The median P value from 10 random sets was used (Extended Data Table S7-1). (****, p<0.0001, ***, p<0.001, **, p<0.01, *, p<0.05).

The difference between the mRNAs resistant to run off and the FMRP targets in these analyses is surprising as over 60% of the mRNAs resistant to run off are also FMRP targets. We examined the mRNAs resistant to run-off separately based on whether or not they were FMRP targets (Extended Data Tables 6-1, 7-1 and Fig S7-1). This revealed that the lack of an effect of FMRP loss in this group was due to the absence of an effect on mRNAs resistant to run-off that were not FMRP targets (Fig. S7-1). Even more striking was the comparison between the occupancy of FMRP targets that were or were not also mRNAs resistant to run-off. FMRP targets that were not also resistant to run-off (85% of this group) showed no increase in occupancy (Fig S7-1). Thus, FMRP targets showed increased abundance and enrichment compared to total mRNAs and decreases in all features in the absence of FMRP (Fig S7-1). In contrast, mRNAs resistant to run-off showed higher occupancy than total mRNAs or FMRP targets and showed decreased abundance and enrichment in the absence of FMRP, but their occupancy was not affected by the loss of FMRP (Extended Data Tables 6-1, 7-1 and Fig. S7-1).

### Peak Analysis of RPFs in the WT and FMR1- Pellet

The distribution of RPFs in stalled ribosomes is dominated by peaks, presumably representing stalling sites. Our previous result showed that RPFs in the peaks from the pellet were enriched with FMRP related mRNA motifs (Anadolu et al., 2023). Thus, to inquire if the loss of FMRP altered the location of stalled ribosomes, peaks were identified in each of the six pellet preparations (3 WT and 3 FMR1-) (Fig. 8A)

**Figure 8:**
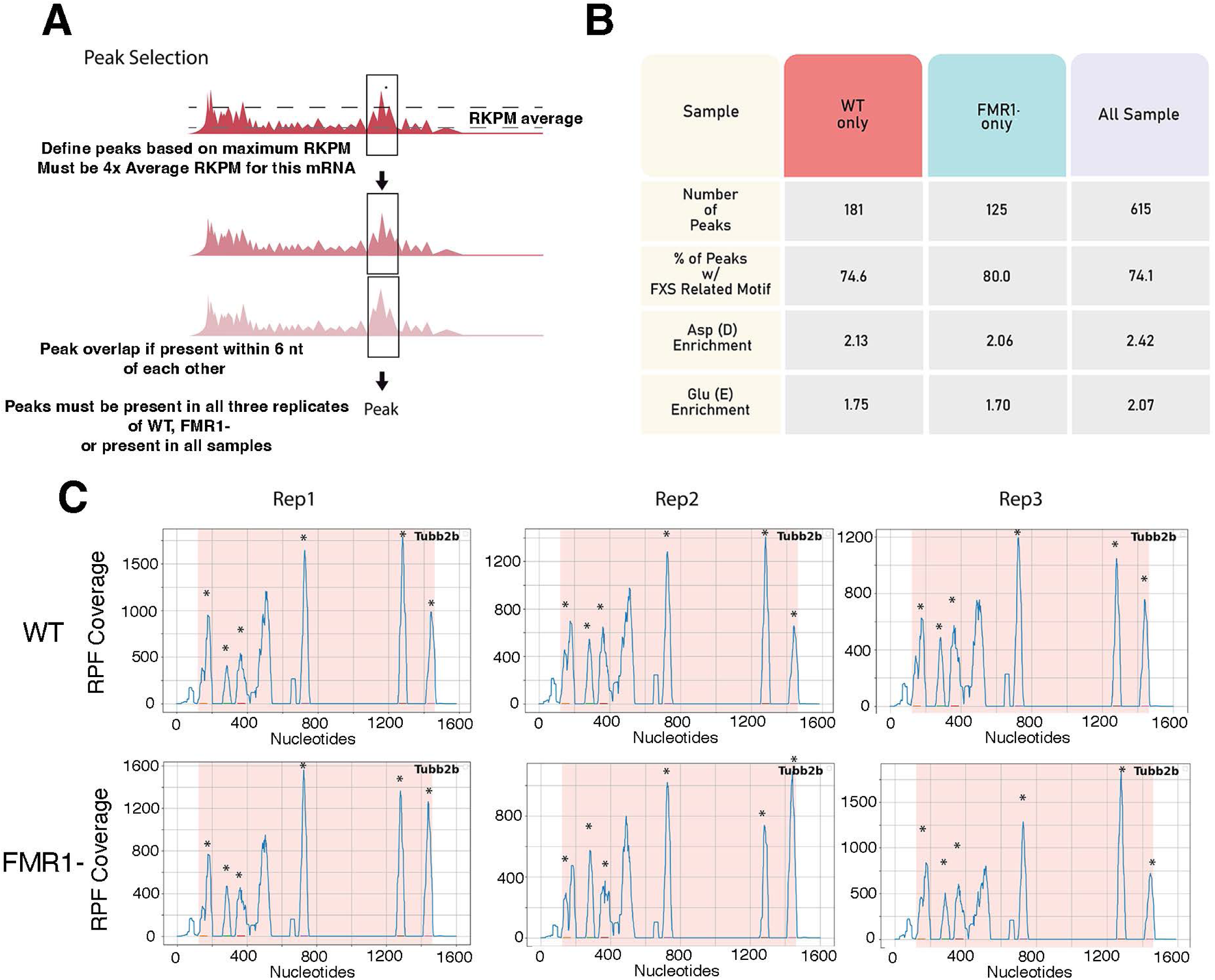
Comparison of RPF peaks in the pellet of WT and FMR1- mice. A) Representation of how peaks of RPFs are selected. B) Table of the number of peaks between replicates of WT Pellet (N = 3), FMR1- Pellet (N = 3) and combined (N = 6), the percentage of peaks with FXS related motif and the enrichment of Aspartate (Asp) and Glutamate (Glu) in the peaks compared to non-peaks (Extended Table S8-2). C) RPF coverage of Tubb2b for the three replicates of WT and FMR1-. Shaded region is the open reading frame. Asterisks indicate consensus peaks (seen in all six samples with peaks within 6 bp).

We identified 615 peaks that were present in all six samples. In contrast, there were many fewer peaks solely present in WT or solely FMR1-: 181 peaks for WT and 125 peaks for FMR1-(Extended Data Table 8-1; Fig. 8B). Thus, most peaks were present in both the WT and FMR1-pellets. In general, the distribution of RPF peaks is quite similar in different biological preparations, as exemplified by the Tubb2b mRNAs (Fig. 8C), a cytoskeletal mRNA that shows high ribosomal occupancy in our samples and that was also examined in the previous manuscript (Anadolu et al., 2023).

Moreover, similar to our previous results, the majority of peaks (74-80%) contain a FMRP related motif, and this was not different in the peaks that were enriched in all the samples and the peaks that were only identified in the WT (i.e not seen in FMR1- mice) or only in FMR1-mice (i.e. not seen in WT mice) (Figure 8B). Previously, we found that these peaks were enriched in negatively charged amino acids, glutamic acid and aspartic acid. This was also true for the peaks here, regardless of whether they were present or absent in the FMR1- mice (Fig. 8B).

### Loss of FMRP Decreases the Number of RNA granules containing Stalled Ribosomes

Since FMRP did not appear to affect the location on mRNAs where the ribosomes were stalled, we next examined the number and size of neuronal RNA granules containing stalled ribosomes in WT and FMR1- hippocampal cultures (Fig. 9A). FMRP is localized to neuronal RNA granules in hippocampal neuronal cultures (Graber et al., 2013; El Fatimy et al., 2016). We examine neuronal RNA granules containing stalled ribosomes in hippocampal neurons using a technique called ribopuromycylation (RPM)(David et al., 2013). The ribosomes covalently link puromycin to nascent peptide chains on ribosomes in a process called puromycylation and these ribosomes are then identified with an antibody to puromycin. Stalled ribosomes are identified by incubating with an initiation inhibitor, in this case homoharringtonine (HHT), to run off translating ribosomes before puromycylation. While in non-neuronal cells, puromycylated peptides leave the ribosome (Enam et al., 2020; Hobson et al., 2020), puromycylated nascent chains are retained on neuronal stalled ribosomes, perhaps due to a peptide-mediated ribosomal stall (Anadolu et al., 2024). Thus, RPM remains appropriate for localizing stalled ribosomes in neurons.

**Figure 9.**
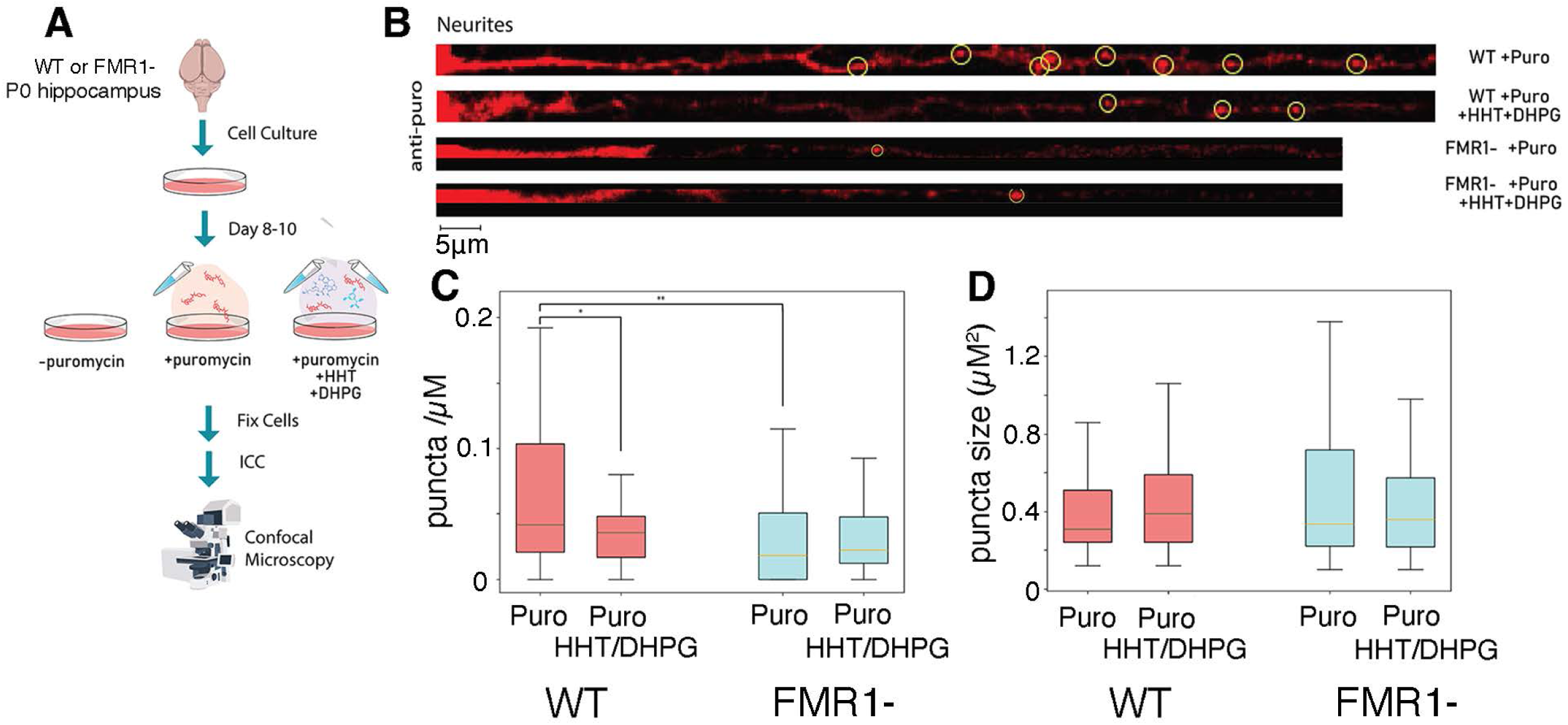
RPM of hippocampal cultures derived from WT and FMR1- mice. A) Summary of the protocol for puromycylation HHT-Runoff and DHPG Reactivation on WT and FMR1- hippocampal culture. B) Representative confocal images for puromycylated ribosomes with or without HHT runoff and DHPG reactivation. Circle denotes puromycin puncta. No visible staining was seen in the absence of puromycin. Scale bar shown below. C) Quantification of RPM puncta density. Numbers are neurites/cultures. WT (42/5); WT DHPG (54/5), FMR1- (41/4), FMR1- DHPG (25/3). One way ANOVA F(3,158)= 5.32, p< 0.005) *, p<0.05 post-Hoc Tukey HSD test. Box and Whisker plot with line representing the median. D) Quantification of size of RPM puncta. WT 189/5; WT DHPG 171/5, FMR1- (118/4), FMR1-DHPG (48/3). Box and Whisker plot with line representing the median. One way ANOVA showed no significance (P>0.05).

We examined RPM puncta (Fig. 9A) in neurites >50 microns from the cell soma as there was too much RPM signal proximal to this to distinguish puncta clearly. We found that the number of RPM puncta were significantly decreased in hippocampal cultures from mice lackingFMRP (Fig, 9B quantified in Fig. 9C). There were no differences in the average size of the remaining puncta (Fig. 9B, quantified in Fig. 9D) and the RPM puncta were equally resistant to both run-off with HHT and competition with anisomycin (Fig. S9-1) consistent with these puncta representing large collections of neuronal stalled ribosomes. Previously we showed that dihydroxyphenylglycine (DHPG), an agonist of metabotropic glutamate receptors (mGluRs) can reduce the number of RPM puncta in these cultures (Graber et al., 2013). The decrease of these RPM puncta is coincident with the reactivation of translation from stalled ribosomes by DHPG at this time point (DIV8-10) (Graber et al., 2013; Graber et al., 2017). Moreover, long-term depression (LTD) induced by DHPG is blocked by elongation inhibitors, but not initiation inhibitors, consistent with DHPG activating protein synthesis required for mGluR-LTD by reactivating stalled ribosomes (Graber et al., 2013; Graber et al., 2017). We replicated this result; DHPG significantly decreased the number of RPM puncta in WT cultures. However, DHPG had no significant effect on the decreased number of puncta in cultures from FMRP- mice (Fig. 9B,C). This is consistent with a loss of DHPG-sensitive RPM puncta in the absence of FMRP.

## Discussion

Overall, there appeared to be no major changes in the biochemical composition of stalled ribosomes in the absence of FMRP (Fig. 1) or the state of the stalled ribosomes (Fig. 2). Most importantly, the places where ribosomes stall (peaks) are not changed in the absence of FMRP since the majority of peaks were present in both WT and FMR1- mice, and the RPFs from FMR1- brains still contained motifs in mRNAs previously shown to associate with FMRP (Fig. 8). Thus, the major conclusion of our study is that FMRP does not affect the locations at which neuronal ribosomes stall.

We did not observe major changes in a DEG analysis of RNA-seq between control and FMR1- mice in P5 brains, although some changes were observed using GSEA. This differs from many studies that do show differences in RNA-seq between these two groups at other stages of development or examining specific brain regions (Thomson et al., 2017; Sawicka et al., 2019; Kurosaki et al., 2022; Ntoulas et al., 2024). It is likely that our use of whole brains, a specific time point, and a relatively small number of biological replicates decreased our ability to see FMRP-related changes in mRNAs that are observed at a more cell-type-specific level and at different developmental times.

In the GSEA analysis of RPFs in the pellet (abundance) there were a number of gene sets that were decreased in the absence of FMRP. These gene sets were enriched in FMRP targets (Fig. 5). Indeed, if we focused just on the FMRP targets, there was a small but highly significant decrease in the relative abundance, occupancy and enrichment of these mRNAs in the absence of FMRP. Interestingly, the mRNAs identified in FMRP Clips could be divided into two classes based on whether they were also identified as mRNAs resistant to run-off. Only the FMRP targets that were also identified as mRNAs resistant to run-off showed increased occupancy compared to other mRNAs. mRNAs resistant to run-off that were not FMRP targets were still significantly decreased in abundance and enrichment, but unlike FMRP targets, loss of FMRP did not affect their occupancy. These results and the finding that peaks of RPFs are not affected by the loss of FMRP are consistent with FMRP not directly regulating stalling, but rather affecting whether stalled mRNAs are found in the pellet. This may be partly due to the role of FMRP in regulating the initiation of FMRP target mRNAs as only initiated mRNAs can be stalled. Indeed, there is considerable evidence that FMRP regulates initiation of a subset of mRNAs (Santini and Klann, 2014; Bagni and Zukin, 2019).

In contrast to the examination of the number of ribosomes and RBPs present in the ribosome clusters in the pellet, we did find a decrease in RPM puncta in hippocampal cultures and the remaining puncta were insensitive to DHPG (Fig. 9). FMRP contains a low complexity domain that allows phase separation to form spontaneously (Zhang et al., 2022). FMRP is subject to post translational modification by neuronal stimuli, such as mGluR-LTD (Niere et al., 2012; Khayachi et al., 2018) that reactivates translation from stalled ribosomes (Graber et al., 2013; Graber et al., 2017). Loss of FMRP leads to an increase in the amount of mGluR-LTD and removes the protein synthesis requirement for mGluR-LTD (Hou et al., 2006; Nosyreva and Huber, 2006). One possibility is that FMRP plays a role in retaining stalled ribosomes within RNA granules by maintaining their liquid-liquid phase, while post-translational modifications of FMRP participates in the controlled disassembly of these granules, enabling translational reactivation. In the absence of FMRP, premature disassembly of RNA granules may lead to premature production of proteins from mRNAs stalled in neuronal RNA granules.

The pellet and the RPM labeled granules are linked by the fact that both the ribosomes in the isolated clusters and in RPM puncta in hippocampal neurites can be puromycylated without dissociation of the puromycylated peptide, and in both puromycylation is less resistant to anisomycin (Graber et al, 2013; Anadolu et al, 2024; Fig. S9-1). However, while stalled ribosomes are enriched in the pellet, it is not clear that only stalled ribosomes from neuronal RNA granules are present in the pellet. Moreover, it is not clear that all stalled ribosomes are sedimented in the pellet. FMRP may be more important in stabilizing ribosome clusters in RNA granules than for forming them. A better understanding of how ribosome clusters move in and out of RNA granules is required to address this issue in more detail.

Our previous results showed a larger protected fragment in RPFs from the pellet, but here we showed that this was due to incomplete nuclease protection. This incomplete digest was much more obvious at the entry site of the ribosome, than at the exit site, as the difference is cleavage is mainly at the 3’end of the protected mRNA (Anadolu et al., 2023)(Fig. 3). It is still unclear whether this is due to increased nuclease resistance of the mRNA near the entrance site of the ribosome due to secondary structures or just differences in nuclease accessibility to the region near the entrance site that is not neuronal specific. Further research on the mechanism of stalling may make this clearer.

Ribosomes containing secretory mRNA are stalled by the signal recognition protein and then co-translationally inserted into the ER. There is evidence that ribosomes attached to the ER can be transported in neuronal processes (Carter et al., 2020; Martin-Solana et al., 2024). Some local mRNAs hitchhike with mitochondria or endosomes (Fernandopulle et al., 2021). Differences in RNA composition between fraction 5/6 and the pellet may be due to increased sedimentation of these other forms of neuronal transport. Indeed, ribophorin, a rough ER protein that associates with ribosomes is enriched in the pellet compared to fraction 5/6 (Fig. 1). However, examination of our standard EM pictures show few examples of ribosomes attached to membranes (Carter et al., 2020; Martin-Solana et al., 2024) and the clustered ribosomes that make up most of the pellet and fraction 5/6 (Fig. 1) do not resemble ribosomes in transit in these other structures.

A major caveat of the present study is that the biochemical studies are only done at a single developmental point (P5 brains) and similarly, the hippocampal neurite studies use a single comparable time point. P5 brains were chosen due to their relative enrichment in neurons (Bandeira et al., 2009) and their lack of myelin. Myelin interferes with the sedimentation of ribosomal clusters (Khandjian et al., 2004; El Fatimy et al., 2016)and the detergents required to remove this interference also interfere with the sedimentation of ribosome clusters (Khandjian et al., 2004; El Fatimy et al., 2016) making comparable studies at later developmental time points not experimentally feasible at the present time. Thus, whether FMRP may play a more important role in determining stalling positions at later developmental time points cannot be ruled out.

## Conclusions

We previously showed that the peaks of RPFs in ribosomes in the pellet contain sequences enriched in FMRP CLIP experiments (Anadolu et al., 2023). Here we show that this is independent of FMRP since the enrichment of these sequences is observed even when FMRP is lost. This suggests that, rather than FMRP driving the recognition of stalling sites, the sequences direct ribosome stalling, and that FMRP associates with stalled ribosomes. Moreover, FMRP does not affect anisomycin-resistant puromycylation of stalled ribosomes, nor does it alter the proteins associated with stalled ribosomes. There is a small decrease in the relative abundance, ribosome occupancy, and enrichment in the pellet of the FMRP clipped mRNAs (FMRP targets) suggesting that these mRNAs do have some specific relationship to FMRP that is independent of FMRP stalling these mRNAs. FMRP also plays a role in RNA granules containing stalled ribosomes since their number is decreased in the absence of FMRP, and the remaining granules are not reactivated by DHPG. We propose that FMRP is important for the regulated dissociation of the liquid-liquid phase neuronal RNA granules containing stalled ribosomes.

## Methods

### Purification of the RNA Granule-Enriched Fraction

All preparations used brains that were flash frozen using either liquid nitrogen or ethanol dry ice bath after dissection. Either three livers, five rat brains (Sprague Dawley; Charles River Laboratory) or 10 mouse brains (C57Bl/6) from WT and FMR1- mouse were used. All animals were 5d old. Samples were homogenized in either the previous (Anadolu et al, 2023) RNA Granule Buffer (20 mM Tris-HCl pH 7.4 (catalog #BP152-1, Thermo Fisher Scientific), 150 mM NaCl (catalog # BP358-212, Thermo Fisher Scientific), 2.5 mM MgCl_2_ (catalog# M33-500, Thermo Fisher Scientific)) or high Mg^2+^ RNA Granule Buffer (20 mM Tris-HCl pH 7.4, 150 mM NaCl, 10 mM MgCl_2_). These buffers were supplemented with 1 mM DTT (catalog #D9163, Sigma-Aldrich), and 1 mM EGTA (catalog # E8145 Sigma-Aldrich) for homogenization. The homogenate was centrifuged 15min in a Thermo Fisher Scientific T865 fixed-angle rotor at 6117 x g at 4 °C to separate debris, such as lipid and extracellular matrix, from the ribosomes. The supernatant was collected with some set aside as starter fraction. The rest of the supernatant was then clarified with 1% IGEPAL CA-630 (catalog #04693132001, Roche) for 5 min at 4 °C Sucrose solution was produced by suspending sucrose (catalog #8550, Calbiochem) with RNA Granule buffer. The samples were loaded onto a 60% sucrose pad in a Sorvall 36 ml tube (Kendro, catalog #3141, Thermo Fisher Scientific) and centrifuged at 56,660 x g for two hours in AH-629 swing-bucket rotor to retrieve the ribosomes. The ribosomes were resuspended in RNA Granule buffer then reloaded onto 15%-60% sucrose gradient and centrifuged at 56,660 x g for 45 min. Each fraction was 3.5 ml and collected from the top.

### Immunoblotting and quantification of enrichment

For immunoblotting, the pellet and fraction 5/6 were ethanol precipitated and resuspended with 1x sample buffer and RNA Granule buffer. The samples were loaded onto 10%, 12% or 15% acrylamide gel according to the observed protein sizes. The gel was either stained with Coomassie Brilliant Blue to look at the protein distribution or transferred onto a 0.45um nitrocellulose membrane (catalog #1620115, Bio-Rad) for immunoblotting. The transferred membranes were stained with Ponceau and imaged. Then, the membranes were blocked with 5% BSA (catalog # A9647, Sigma Aldrich) in Tris-buffered saline with Tween before incubation with primary antibodies— rabbit anti-s6 (1:10,000) (catalog #2217, Cell Signaling Technology), rabbit anti-FMRP (1:500)(catalog #4317, Cell Signaling Technology), rabbit anti-UPF1 (1:10,000)(catalog #ab133564, Abcam), mouse anti-Stau2 (1:1000)(catalog #MM0037-P, MediMabs), anti-eEF2 (1:1000)(catalog #2332S, Cell Signaling Technology), rabbit anti-Pur-alpha (1:1000)(catalog #ab79936, Abcam), rabbit anti-ZBP1 (1:500)(catalog # NBP1-7685 Novus Biologicals, rabbit anti-TIA1 (1-1000)(Catalog #12133-2-AP Proteintech), rabbit anti-hNRPA2B1 (1:1000)(catalog #NB120-6102), mouse monoclona anti-ribophorin (1:200)(Catalog #sc-48367 Santa Cruz Biotechnology) and anti-mouse puromycin (1:1000)(catalog #ab2619605, Developmental Studies Hybridoma Bank). Membranes were washed with TBS-T after incubation. HRP-conjugated secondary antibodies such as anti-rabbit HRP (1:10,000)(catalog #31460, Thermo Fisher Scientific) and anti-mouse HRP (1:10,000)(catalog #31430, Thermo Fisher Scientific) were incubated with the membranes for detection. ECL reaction was performed for imaging, and the images were scanned and quantified by ImageJ software. The single band for each protein was selected and quantified using ImageJ’s Gel analysis Macro. For S6, the levels in the pellet and fraction 5/6 were directly compared based on the fraction of the samples loaded on the gel . For other proteins, the levels in the pellet and fraction 5/6 were first calculated based on the fraction of the samples loaded on the gel and then normalized to the relative S6 levels found in the pellet and fraction 5/6 from this preparation to provide a RBP/ribosome ratio. The enrichment for the pellet was determined by dividing this normalized value between the pellet and fraction 5/6. . A two tailed, one sample t-test was performed between the enrichment and 1 with Bonferroni correction for multiple T-tests to test for significant enrichment. A two tailed, Welch’s t-test was performed between the enrichment of WT and FMR1- with Bonferroni correction for multiple T-tests to observe the differences between the two groups.

UV absorbance

2 µL of each fraction was measured using Nanoblot 1000 Spectrophotomer (Thermo Fisher Scientific). Averages of the value between three independent experiments from WT and FMR1- mice were used.

### Inhibition of Puromycylation by Anisomycin

The liver ribosomes were extracted from P5 Spraque Dawley rats through the identical method as the RNA Granules, but without the last spin since liver does not contain appreciable levels of ribosomes in the pellet. All ribosomal fractions used were incubated for 5 min in 1) RNA Granule buffer 2) 100μM puromycin (catalog# P7255, Sigma Aldrich), or 3) 100μM puromycin and 100μM anisomycin (catalog# A9789, Sigma Aldrich). The samples were then ethanol precipitated, immunoblotted and quantified via the method stated above. The percentage of puromycin resistant to anisomycin inhibition were calculated by dividing anti-puro signal from the puromycin and anisomycin added sample against the anti-puro signal from the puromycylated sample within each replicate.

### Digestion and extraction of the monosomes

Fraction 5/6 samples were loaded onto a 60% sucrose pad and centrifuged to concentrate the samples, while the pellet samples were resuspended from the pellet using 1 ml of normal or high MgCl_2_ Granule buffer. For normal nuclease treatment groups, 1µl of RNAase I (100U/µ l; catalog #AM2294, Thermo Fisher Scientific) was administered to the ribosomes and rotated at 4°C for 30min. Then, 4µl of SuperaseIN (20U/µl; catalog # AM2969, Invitrogen) was added to the solution to halt the reaction. The samples are then loaded onto 15% to 60% sucrose gradient and centrifuged at 56,660 x g for 45 minutes to retrieve the monosomes from fraction 2 and 3. For high nuclease treatment group, RNAse I (10 U/µl, catalog #N6901K, Epicentre) was adjusted to the concentration of ribosomes via the A260 read from the Nanodrop. The OD obtained from the A260 read equals 6 unit. In other words, the amount of nuclease (µl) added equals A260 * 6 (U)/ 10 (µl/U). In addition, the samples are incubated at room temperature for 30 min rather than at 4 °C. Then, 6µl of SuperaseIN were added to halt the reaction. The samples were then spin at 68,000 x g for 3hrs on a Beckman tabletop ultracentrifuge to concentrate the monosomes, which pellets. The RNA from the monosomes was extracted using the trizole chloroform method followed by isopropanol precipitation to concentrate the samples. The samples were loaded onto Urea gel (catalog # EC68852Box, #EC68752Box, #EC62152Box, ThermoFisher) to select for RPF size. Segments between 25b and just above 40b were excised and retrieved to account for the possible longer fragments of the RPFs. The excised gels were frozen on dry ice for 30 minutes and then thawed overnight at room temperature. The RNA was extracted again with trizole chloroform extraction.

#### Linker Ligation

The protocols follow improvements in RPF ligation (McGlincy and Ingolia, 2017). The concentration from the RNA footprint was calculated from Bioanalyzer small RNA kid (catalog # 5067-1548, Agilent). An equal amount of RNA was calculated and transferred to a new tube for each sample to ensure each sample has a relatively equal amount after pooling. 3 different linkers (NI-810, NI-811, BI-812) were attached to each of the replicates. The samples were first dephosphorylated with T4PNK linker (catalog # M0351L, NEB), and then pre-adenylated linkers are attached through T4 Rnl2 (catalog # M0351L, NEB). The linked RNA was purified through excision of urea gel between 50bp and 70bp. Samples with the same variables were then pooled together with their gel fragment combined. The RNA was extracted from the gel with the steps stated previously containing all three replicates, followed by Reverse transcription. MyOne Streptavidin C1 DynaBeads (catalog#65001, ThermoFisher) was used for rRNA depletion. Lastly, PCR was performed. The samples were sequenced through the McGill University Genome Center on NovaSeq S1/2 flow cells.

### RPF and RNA-seq data analysis

The raw reads were processed using Cutadapt (version 2.10), where sequences were trimmed to remove adapters and low-quality bases. Subsequently, demultiplexed each pooled reads runs using UMI barcodes (ID1=ATCGT, ID2=AGCTA and ID3=CGTAA). Additionally, options were used to trim N bases (--trim-n), discard sequences shorter than 18 bases (--minimum-length=18), allow a minimum overlap of 5 bases (--overlap=5), and omit insertions or deletions (--no-indels). Untrimmed reads were discarded, and trimming logs were saved for quality control. Reads mapping to non-coding RNAs (ncRNAs) and ribosomal RNAs (rRNAs) were filtered out using Bowtie2 (version 2.3.5), aligning the reads against rRNA and ncRNA sequences from the Mus musculus reference (GRCm38, Ensembl release 102). The Bowtie2 parameters were set to maximize sensitivity (--very-sensitive). Unmapped reads were further aligned to the Mus musculus genome (GRCm38, Ensembl release 102) using the STAR aligner (version 2.7.8a). The output was generated as transcript coordinate-sorted BAM files (--outSAMtype BAM SortedByCoordinate --quantMode TranscriptomeSAM). PCR duplicates were identified and marked using *Picard MarkDauplicates* (version *2.26.6)*. These duplicates were then removed using *samtools* (version 1.19.2). Further, raw counts were obtained using featurecounts (version 2.2.0).

For RPKM and all DEG analysis, non-protein coding genes were removed. RPKM, reads per kilobase per million mapped reads, were calculated for abundances, which normalized the raw counts against gene length and total count. Occupancy for individual groups were calculated through performing DEG analysis between pellet raw count replicates and whole brain RNA-seq replicates, and enrichment for individual groups were calculated through performing DEG analysis between raw count pellet replicates and raw count fraction 5/6 replicates. For comparison between WT and FMR1- groups, DEG Analysis was calculated as follow: abundances, raw count WT vs raw count FMR1- ; occupancy, raw count WT pellet divided by whole brain RNA-seq vs raw count FMR1- pellet divided by whole brain RNA-seq, enrichment, raw count WT pellet divided by raw count WT fraction 5/6 vs raw count FMR1- pellet divided by FMR1-KO fraction 5/6..

The Differential Expression Gene Analysis was performed via R studio packages in *edgeR* from Bioconductor adapted for RNA-seq (Wu et al., 2021). Data were normalized for library size differences and filtered to remove lowly expressed genes. Results were ranked and evaluated based on adjusted p-value (FDR), which was calculated through the Benhamini-Hochberg method. To visualize sample clustering and data quality, Principal Component Analysis (PCA) was performed on log-transformed counts-per-million (CPM) values using the **stats** and **edgeR** packages low expression gene was filtered and normalized. For GSEA, resulting gene lists were ranked and processed for Gene Set Enrichment Analysis (GSEA) using the fgsea package against MSigDB C5 (GO) mouse gene sets. Pathway enrichment was restricted to sets containing 10–500 genes, with significance prioritized by the Benjamini-Hochberg adjusted p-value. GO Enrichment Analysis was performed with clusterprofiler (Wu et al., 2021) and org.Mm.eg.db from Biomanager using R studio (Yu, 2024). Comparison to data online was performed via jupyter notebook and graphed by *boxplot* in *matplotlib*. The significance of each dataset comparison was done through performing t-test between the overlapped gene in our data and non-overlapped genes through the *statistics* function from python. The p-value was Bonferroni corrected.

We used transcript coordinate Bam files for peak analysis. Regions of the mRNA that have an abundant amount of RPFs were referred to as peaks (Anadolu et al., 2023). The peaks previously were selected through marking the inflection point within the abundances of RPFs. Here, we identify the highest point (zenith) in the peak after mapping RPFs to transcripts. To be identified as a peak, the zenith of an abundance site for the reads must be 4x higher of the average of the total transcript. Moreover, this zenith must be detected in all the biological samples (zenith within 6 nucleotides for each replicate). We first determine total peaks in all six samples (WT and FMR1-) and then determined peaks that were in all WT samples, but not in FMR1- samples (WT peaks; peaks lost in the absence of FMRP) and peaks that were present only in the FMR1- samples, but not in any WT samples (FMRP peaks).

For identification of motifs in the peaks, the zenith was extended on both sides by 17nt. Motif occurrences in the peaks were identified using FIMO from the MEME suite (Bailey et al., 2015) and universalmotif (Tremblay, 2024). Percentage peaks with FXS related motifs were identifying only the positive strand and limiting the score, or log-likelihood ratio, to 5, and normalizing against the total peaks. Biopython was utilized to convert peak sequences to amino acids.

### Cryo-electron microscopy

The purified fractions were treated with nuclease as described (Anadolu et al., 2023) before being applied to the electron microscopy grids. The samples were applied to the grid in buffer containing 20mM Tris-HCl, pH 7.4; 150 mM NaCl; 10 mM MgCl^2+^, and they were applied at a concentration of 180 nM. Cryo-EM grids (c-flat CF-2/2-2C-T) used for these samples were washed in chloroform for two hours and treated with glow discharge in air at 15 mA for 20 seconds. A volume of 3.6 μL was applied to the grid before vitrification in liquid ethane using a Vitrobot Mark IV (Thermo Fisher Scientific Inc.). The Vitrobot parameters used for vitrification were blotting time 3 seconds and a blot force +1. The Vitrobot chamber was set to 25 °C and 100% relative humidity.

All datasets were collected at FEMR-McGill using a Titan Krios microscope operated at 300 kV and equipped with a Gatan BioQuantum LS K3 direct electron detector. The software used for data collection was SerialEM (Schorb et al., 2019). Images were collected in counting mode at a nominal magnification of 81,000x, producing images with a calibrated pixel size of 1.09 Å. Movies for all datasets were collected using 30 frames with a total dose of 40 e^−^/Å^2^.

### Image processing

All the image processing steps were done using cryoSPARC v4.5 software (Punjani et al., 2017). A total of 10,866 cryo-EM movies were corrected for beam-induced motion correction using Patch Motion Correction using default settings that included using information up to 5Å resolution when aligning frames, a B-factor of 500 and a 0.5 calibrated smoothing constant applied to the trajectories. CTF parameter estimation was done using Patch CTF estimation with default settings. The minimum and maximum resolution considered to estimate the CTF parameters were 25 Å and 4 Å, and the minimum and maximum defocus values were set up at 1,000 and 50,000 Å. The corrected micrographs were then curated by Manually Curate Exposures. For the particle-picking step, the Blob Picker program was first applied to 2,000 obtained micrographs using a circular blob with a minimum and maximum particle diameter of 200 Å and 480 Å. The maximum resolution considered during picking was 20 Å. The angular sampling used was 5 degrees, and the minimum particle separation distance was 0.5 (in units of particle diameters). The picked particles were extracted with a box size of 448 pixels, which was reduced to 112 pixels, and then subjected to 2D classification to generate the 2D templates for subsequent Template Picker using the 2,000 micrographs. In the 2D Classification step, we requested 50 classes and we selected 0.85 and 0.99 as inner and outer window radius. The maximum resolution considered in the images was 9 Å, and we used an initial uncertainty factor of 2. The remaining settings were used with their default parameters. The particles obtained from the Template Picker were curated again by 2D classification to remove the bad particles using the same job parameters. The curated particles were selected and used to train a model using Topaz (Bepler et al., 2019) with default settings, including a minibatch size of 128 and an expected number of particles of 165 per micrograph. The trained model was then used to pick particles from all the micrographs. The selected particles underwent 2 cycles of 2D Classification to remove junk particles. The obtained particles were then subjected to the subsequent particle curation step, which combined Ab-initio Reconstruction and Heterogeneous Refinement programs using default settings. For the Ab-Initio step, we selected 0.85 and 0.99 as inner and outer window radius, requested 3 classes and a maximum and minimum resolution to consider of 35 Å and 12 Å. All other parameters for this routine were used with the default settings and values. The three initial Ab-initio models generated were subsequently used in a Heterogeneous Refinement using default parameters to separate the particles into multiple classes. The particles assigned to the class with unidentifiable features were discarded, whereas the particles assigned to reconstructions with ribosomal features were merged for subsequent processing. The total number of good particles after particle curation was 890,644.

To explore the structural heterogeneity, the curated set of particles was used to generate a consensus map with Non-Uniform Refinement at default settings and C1 symmetry. The aligned particles were then subjected to 3D Variability Analysis requesting 3 orthogonal principal modes and the subsequent 3D Variability Display 3D job was ru in cluster mode with all settings at default values. The number of clusters requested across experiments, ranging from 3 to 5.

Results were filtered at 9 Å. Overall, we performed two rounds of the 3D Variability Analysis combined with 3D Variability Display to address sample heterogeneity. Resulting maps from the exhaustive 3D classification were visually inspected in (Pettersen et al., 2004; Pettersen et al., 2021) and groups of particles representing similar structural features were merged. To obtain high-resolution structures, the particles from each class were extracted with a box size of 448 pixels and refined with Non-Uniform Refinement. The refinement jobs were run under default settings with C1 symmetry, optimized per-particle defocus, optimized per-exposure group CTF parameters and options ‘Fit Spherical Aberration’, ‘Fit tetrafoil’ and ‘Fit anisotropic Magnification’ activated. To improve local resolution, Local Refinement was performed under default settings to refine the two subunits (40S and 60S) independently for all the classes. Particle Subtraction run under default settings was used before Local Refinement to subtract the signal from the particle stacks that will not be used for Local Refinement. Composite maps were obtained by aligning them to the consensus maps of the 80S ribosome and merging the 60S and 40S cryo-EM maps from local refinement using the ‘vop add’ command in Chimera.

Average resolution estimation and local resolution analysis were done with cryoSPARC using the gold-standard approach (Henderson et al., 2012). For each map, we calculated the following FSC plots: ‘No mask’: the raw FSC calculated between two independent half-maps reconstructed from the data and no masking applied. ‘Loose’: FSC calculated after applying a loose soft solvent mask to both half maps. The loose mask is calculated by thresholding the density map at 50% of the maximum density value. The resulting volume is dilated to create a soft mask. Voxels in the mask within 25 angstroms of the thresholded region receive a mask value of 1.0. Voxels between 25 and 40 angstroms fall off with a soft cosine edge, and voxels outside 40 angstroms receive a value of 0.0. ‘Tight’: FSC calculated after applying a tight soft solvent mask to both half maps. The tight mask is calculated by the same procedure as the loose mask, except that the dilation distances are 6 angstroms for the value 1.0 distance and 12 angstroms for the value 0.0 distance. ‘Corrected’: FSC curve calculated using the tight mask with correction by noise substitution (Chen et al., 2103). In brief, the two half maps have their phases randomized beyond a certain resolution, then the tight mask is applied to both, and an FSC is calculated. This FSC is used along with the original FSC before phase randomization to compute the corrected FSC. This accounts for correlation effects induced by masking. The resolution at which phase randomization begins is the resolution at which the no-mask FSC drops below the FSC = 0.143 criterion. Cryo-EM map visualization was performed in UCSF Chimera and Chimera X (Pettersen et al., 2004; Pettersen et al., 2021).

### Ribopuromycylation Assay

WT mouse and FMR1- mouse hippocampal neurons were dissected from P0 mouse and cultured on to poly-L-lysine (PLL)-coated 18mm coverslips as previously described (Langille et al., 2019). The hippocampal neurons were incubated with Neurobasal media supplemented with 1% (vol/vol) N2 and penicillin/streptomycin, 2%(vol/vol) B27 and 0.5 mM GlutaMAX (Life Technologies). Treatment groups: +puro (puromycin), -puro, +puro + anisomycin, +puro +HHT (homoharringtonine), and +puro + HHT + DHPG (S)-3,5-Dihydroxyphenylglycine) were assigned to cells and added to the culture on day 8. Cells that were assigned to HHT or HHT + DHPG conditions were preincubated with supplemented neural basal media as stated above with 5µM HHT (catalog # 1416, Tocris Bioscience) or with 5µM HHT and 100µM DHPG (catalog # 0805, Tocris Bioscience) for 15min in 37°C to ensure sufficient time for ribosomes to runoff and for DHPG to reactivate stalled ribosomes. The control +puro, - puro, and + puro +Aniso groups were incubated with supplemented neural basal media. The solution was then removed from all groups and replaced with new supplemented neural basal medium for the control group and supplemented neural basal medium with 100µM Puromycin or 100µM Puro and 100µM Anisomycin. All groups were incubated at 37 °C for 5min similar to previous experiments (Graber et al, 2013; Langille et al 2019). The cultures were then placed on ice and washed with HBS supplemented with 0.0003% digitonin for 2min, followed by 3x wash with HBS. The cultures were then placed at room temperature and fixed with 4% paraformaldehyde for 30 min. Upon completion, the cell cultures were washed with PBS three times, sealed in the 1x PBS, and placed at 4°C till immunocytochemistry.

### Immunocytochemistry

Cultures were treated with 0.1% Triton-X 100 with 30% sucrose in PBS for 10min to allow for permeabilization of cell membrane followed by 15min of quench solution (55mM ammonium chloride in 1x PBS). The cultures were then washed in 1xPBX for three times before blocking with 1% BSA in PBS for 30min. The cultures were then incubated with primary antibody solution (1:1000 mouse anti-puromycin (DSHB Hybridoma Product PMY-2A4)) in 1% BSA and 1xPBS for 1hour. The primary antibody solution was then removed, and the cultures were then washed 3 times with 1xPBS. Secondary antibody solution (anti-mouse 1:1000 Alexa Fluor 745) was then added to the cultures and incubated for 1hour. The secondary antibody solutions were then removed, and the cultures were washed 3 times with 1xPBS. The culture-containing coverslips were then removed and mounted using 10µL of Dako mounting medium. The coverslips were stored in the dark before imaging.

### Confocal Microscopy

Cells were imaged using a Zeiss LSM-900 confocal microscope with a 63x oil immersion objective. The images were then assigned numbers and randomized to lower potential biases. ImageJ was used to straighten the neurites.

### Quantification of RPM

Only cells with straightened neurites over 75 μm were used. The person doing the quantification was blind to the treatment or genotype. The images were converted to 8 bits and then thresholded at a minimum of 180/256 pixel intensity although there was a manual component to this quantification based on the relative brightness of the image. The analyze particle Macro of Image J was used to identify puncta with size criteria of 0.15 to 2 μm^2^ and circularity of 0.4-1.0. Only puncta>50 μm from the soma were considered for this analysis.

## Data Availability

The cryo-EM maps obtained in this study have been deposited in the Electron Microscopy Data Bank (EMDB), and the accession codes are detailed in Supplemental Table 4-1. All sequences are available in the GEO database (GSE291701).

## List of Supplemental Information

### Supplemental Extended Data Tables

Extended Data Table S3-1. List of Genes with high ratios of long reads/short reads in WT Pellet RPFs

Extended Data Table S5-1. RNA-Seq and RKPM for all samples for all genes. Occupancy and Enrichment calculations for WT and FMR1- samples.

Extended Data Table S5-2. DEG analyses comparing WT and FMR1-

Extended Data Table S5-3 GO analysis of WT and FMR1-

Extended Data Table S5-4 GSEA analysis of differences between WT and FMR1- including Percentage of mRNAs matched that are FMRP targets.

Extended Data Table S6-1 Abundance, Occupancy and Enrichment for WT and FMR1- samples matching the Shah, Maurin and Darnell Lists and subsets of these lists.

Extended Data Table S7-1. Abundance, Occupancy and Enrichment comparisons of the ratio between WT and FMR1- for mRNAs matching the Shah, Maurin and Darnell Lists and subsets of these lists.

Extended Data Table S8-1. List of all Peaks as well as Motif counts.

Extended Data Table S8-2 Amino acid enrichment of Peaks

## Supplemental Figures

**Figure S3-1.**
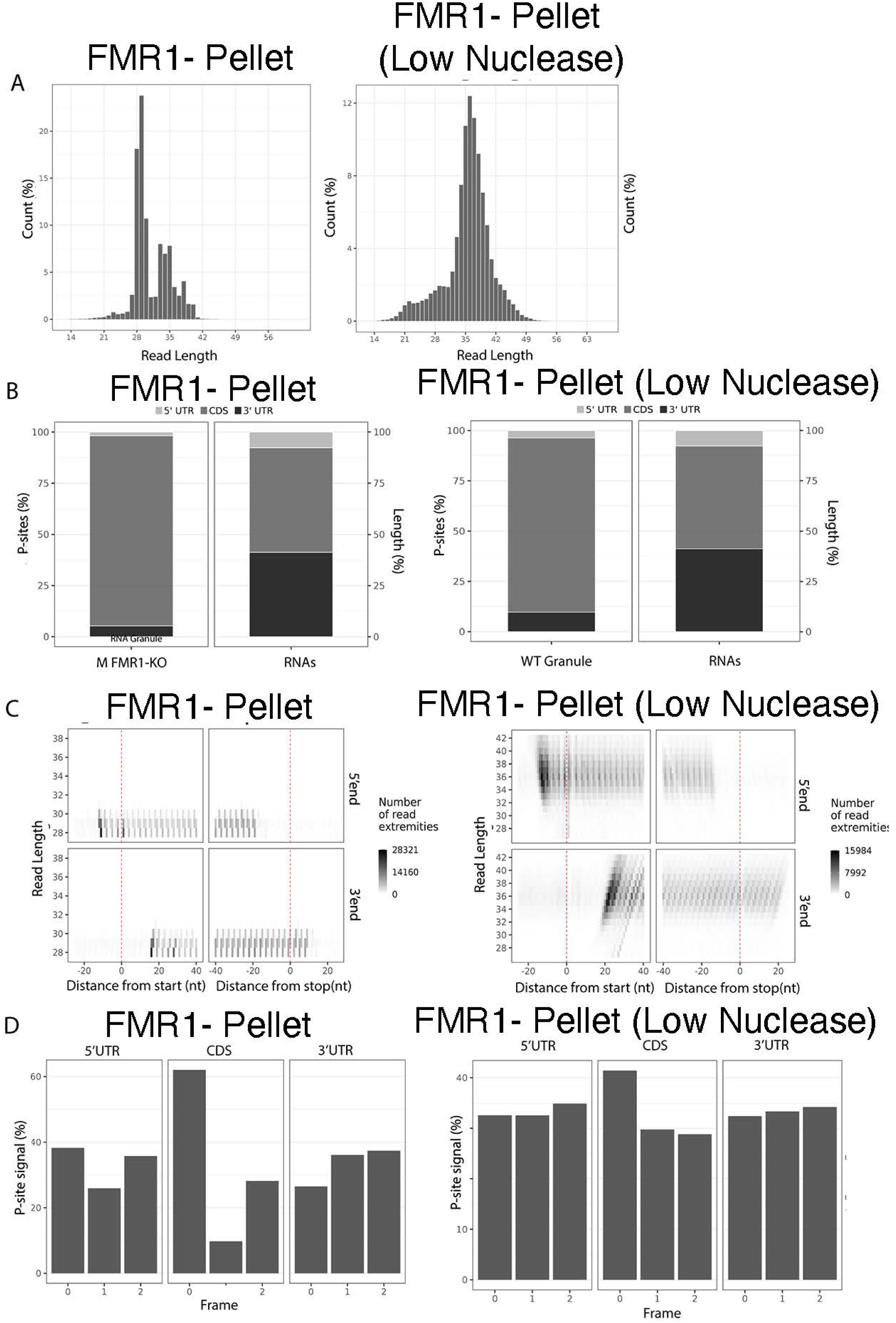
Higher Nuclease reduces size of RPFs in FMR1- pellet . A) Size distribution of normalized footprint reads from the FMR1- Pellet fraction under standard or low nuclease treatment. B) Representative image for read coverage for the FMR1- Pellet fraction either with standard or low nuclease treatment, UTR, untranslated region; CDS, coding sequence for both RPFs and RNA-seq libraries. C) Representative image for the number of read extremities (shading) for each read length (Y-axis) based on the distance from start(left) to stop(right) with the 5’ end (top) and 3’ end (bottom) for the FMR1- pellet fraction with either standard or low nuclease treatment. E) Representative image for the periodicity statistics for RPFs in different regions of the mRNA for standard or low nuclease treatment. Although the representative images above included only one replicate, similar results were observed across all three replicates.

**Figure S3-2.**
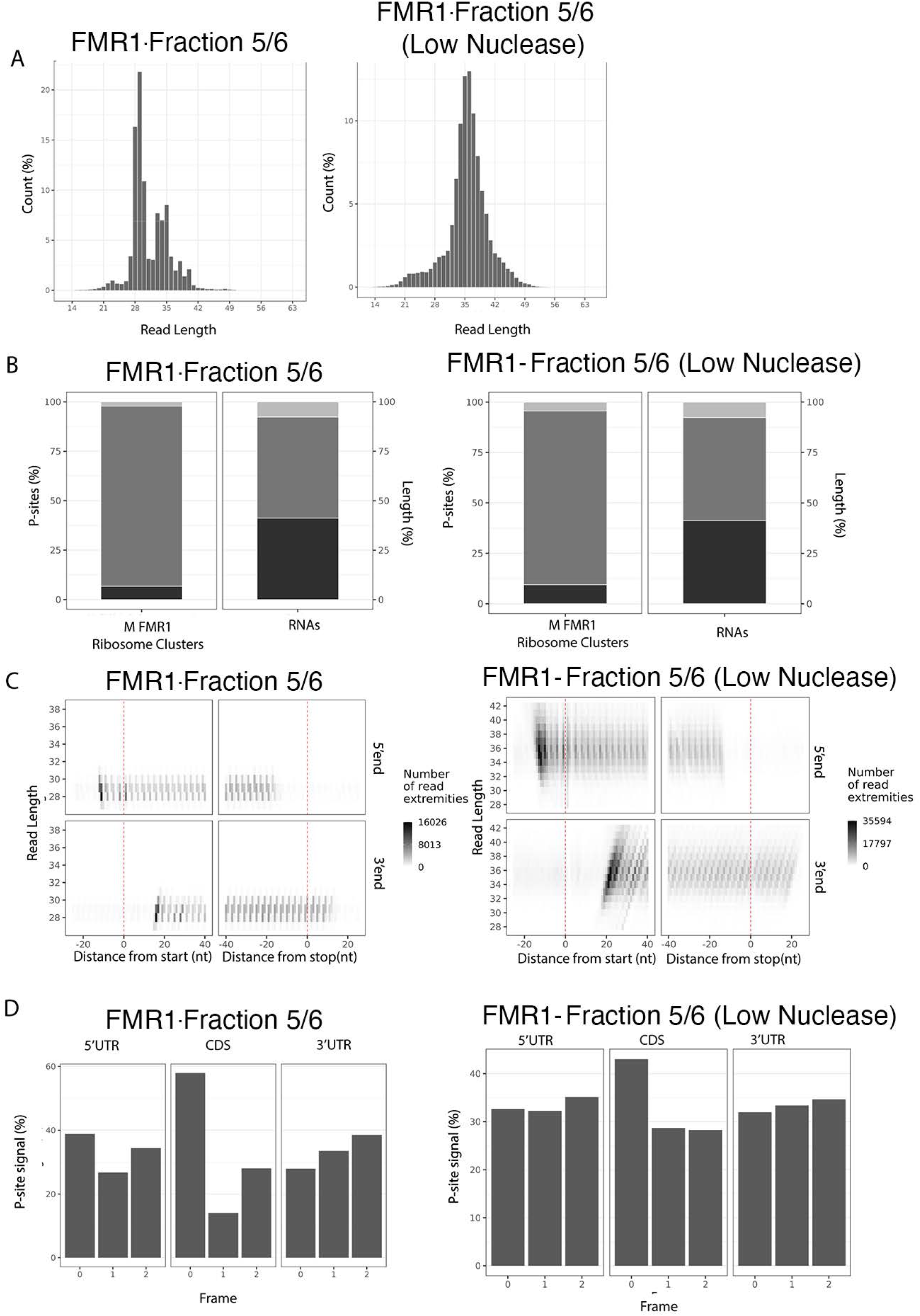
Higher Nuclease reduces size of RPFs in FMR1- Fraction 5/6. A) Size distribution of normalized footprint reads from the FMR1- fraction 5/6 under standard or low nuclease treatment. B) Representative image for read coverage for the FMR1- fraction 5/6 either with standard or low nuclease treatment, UTR, untranslated region; CDS, coding sequence for both RPFs and RNA-seq libraries. C) Representative image for the number of read extremities (shading) for each read length (Y-axis) based on the distance from start(left) to stop(right) with the 5’ end (top) and 3’ end (bottom) for the FMR1- fraction 5/6 with either standard or low nuclease treatment. E) Representative image for the periodicity statistics for RPFs in different regions of the mRNA for standard or low nuclease treatment. Although the representative images above included only one replicate, similar results were observed across all three replicates.

**Figure S3-3.**
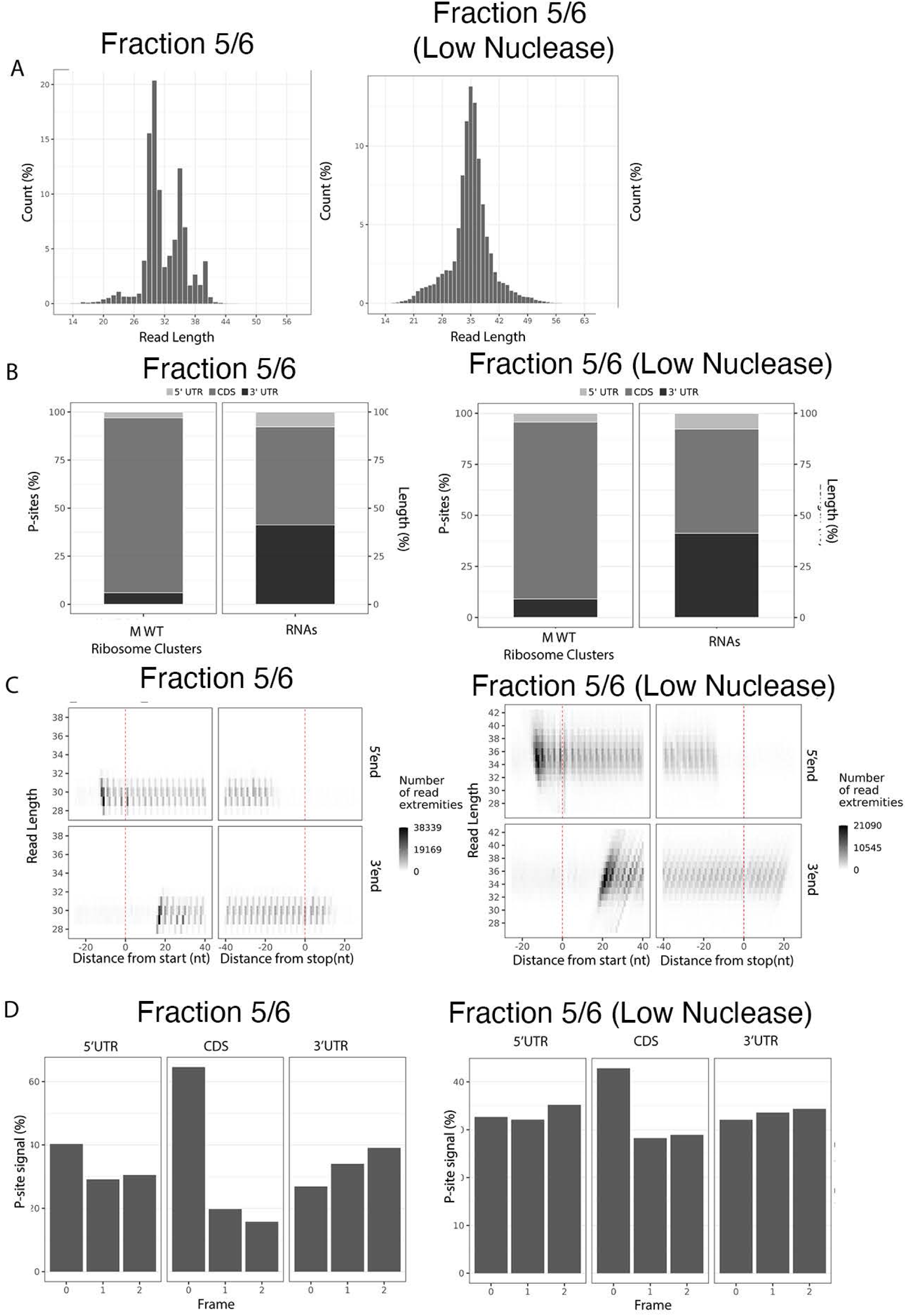
Higher Nuclease reduces size of RPFs in WT fraction 5/6. A) Size distribution of normalized footprint reads from the WT fraction 5/6 under standard or low nuclease treatment. B) Representative image for read coverage for the WT fraction 5/6 either with standard or low nuclease treatment, UTR, untranslated region; CDS, coding sequence for both RPFs and RNA-seq libraries. C) Representative image for the number of read extremities (shading) for each read length (Y-axis) based on the distance from start(left) to stop(right) with the 5’ end (top) and 3’ end (bottom) for the WT fraction 5/6 with either standard or low nuclease treatment. E) Representative image for the periodicity statistics for RPFs in different regions of the mRNA for standard or low nuclease treatment. Although the representative images above included only one replicate, similar results were observed across all three replicates.

**Supplementary Figure S4-1.**
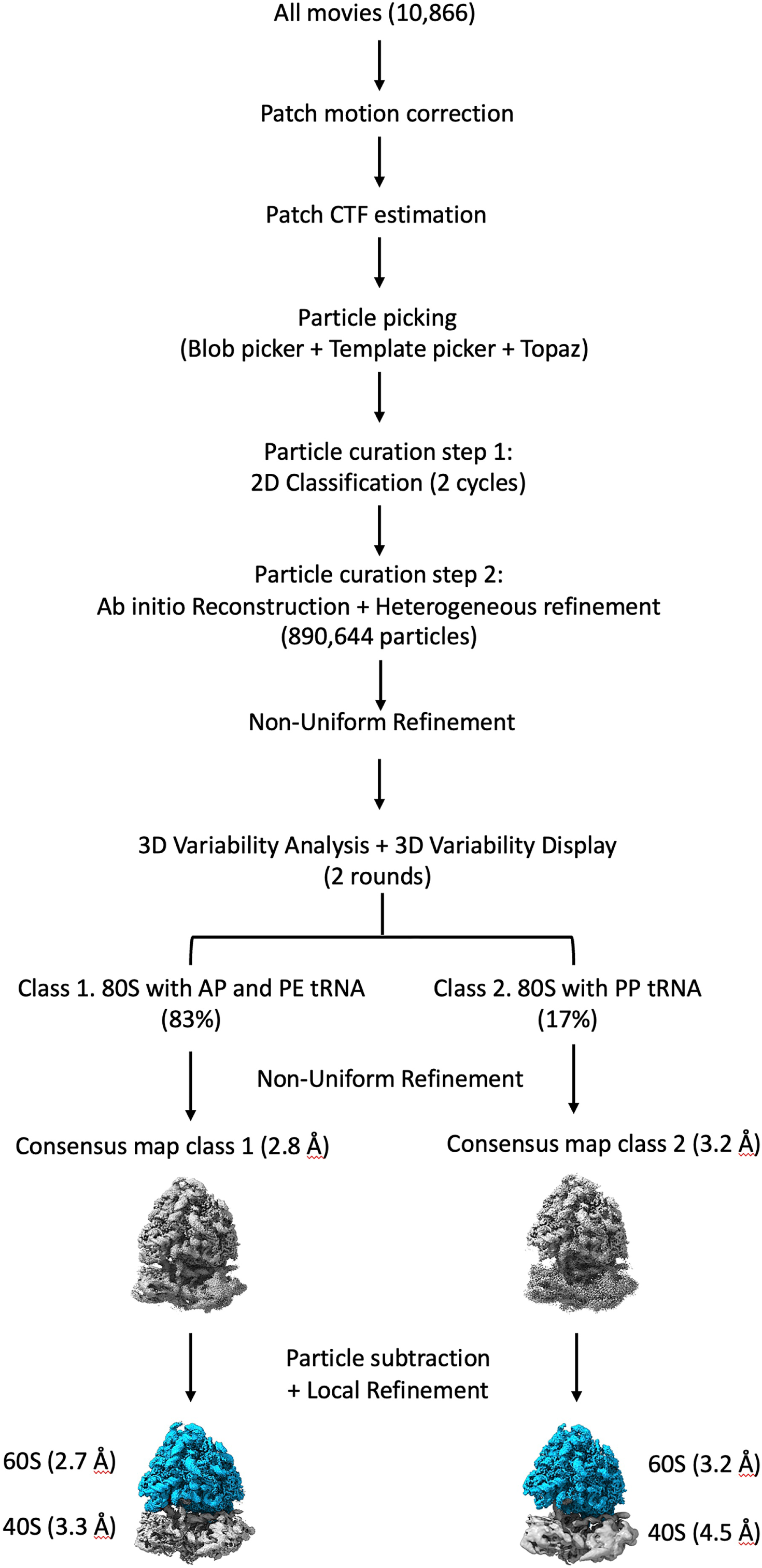
Single particle analysis image processing workflow. processing workflow. The cryo-EM dataset obtained from the pellet fraction after purification in high-magnesium buffer and RNase I treatment was subjected to the image-processing workflow shown in the figure. The diagram shows the main image processing steps performed on this dataset and the two main ribosome populations identified by the image classification approaches. The resolutions for the consensus maps and for each subunit in the maps obtained through local refinement are also indicated.

**Supplementary Figure S4-2.**
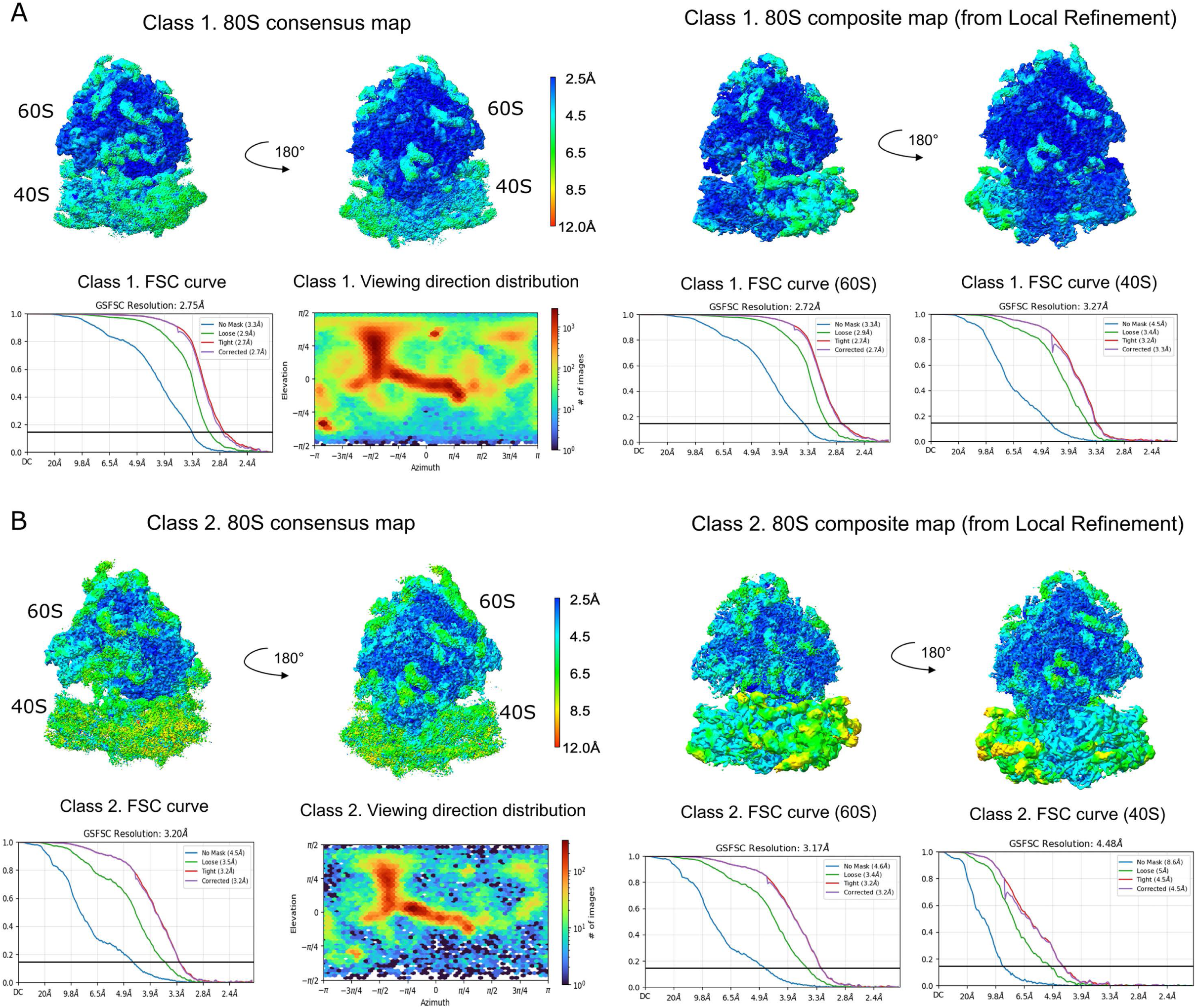
Resolution analysis of the two major classes of 80S ribosomes in the Granule Fraction purified under high magnesium conditions. Consensus cryo-EM maps (left panels) were initially calculated for class 1 (A) and class 2 (B) from the RNase I-treated pellet Fraction purified under high magnesium buffer conditions. These maps were subsequently refined by local refinement by dividing the 80S ribosome into two major bodies, the 60S and the 40S particles. The top panels in (A) and (B) show the local resolution analysis of the 80S consensus and composite maps obtained using local refinement. Maps are colored according to their local resolution using the color coding indicated in the scale bars. The Fourier shell correlation (FSC) curves for the consensus maps and each one of the subunits after the maps were subjected to local refinement are shown for class 1 and 2. For each class, we show the following FSC plots: ‘No mask’, ‘Loose’, ‘Tight’, and ‘Corrected’. These plots were calculated as described in the Methods section. We used a FSC threshold of 0.143 to report the overall resolution of the maps. The ‘Viewing direction distribution’ plot in panels (A) and (B) show the orientation distribution of the particles contributing to the cryo-EM map for each one of the classes. These plots are 2D histograms that show the number of particles with a viewing direction at a particular elevation/azimuth bin. The number of particles can be inferred by the color code scale bar to the right of each plot.

**Supplemental Figure 5-1.**
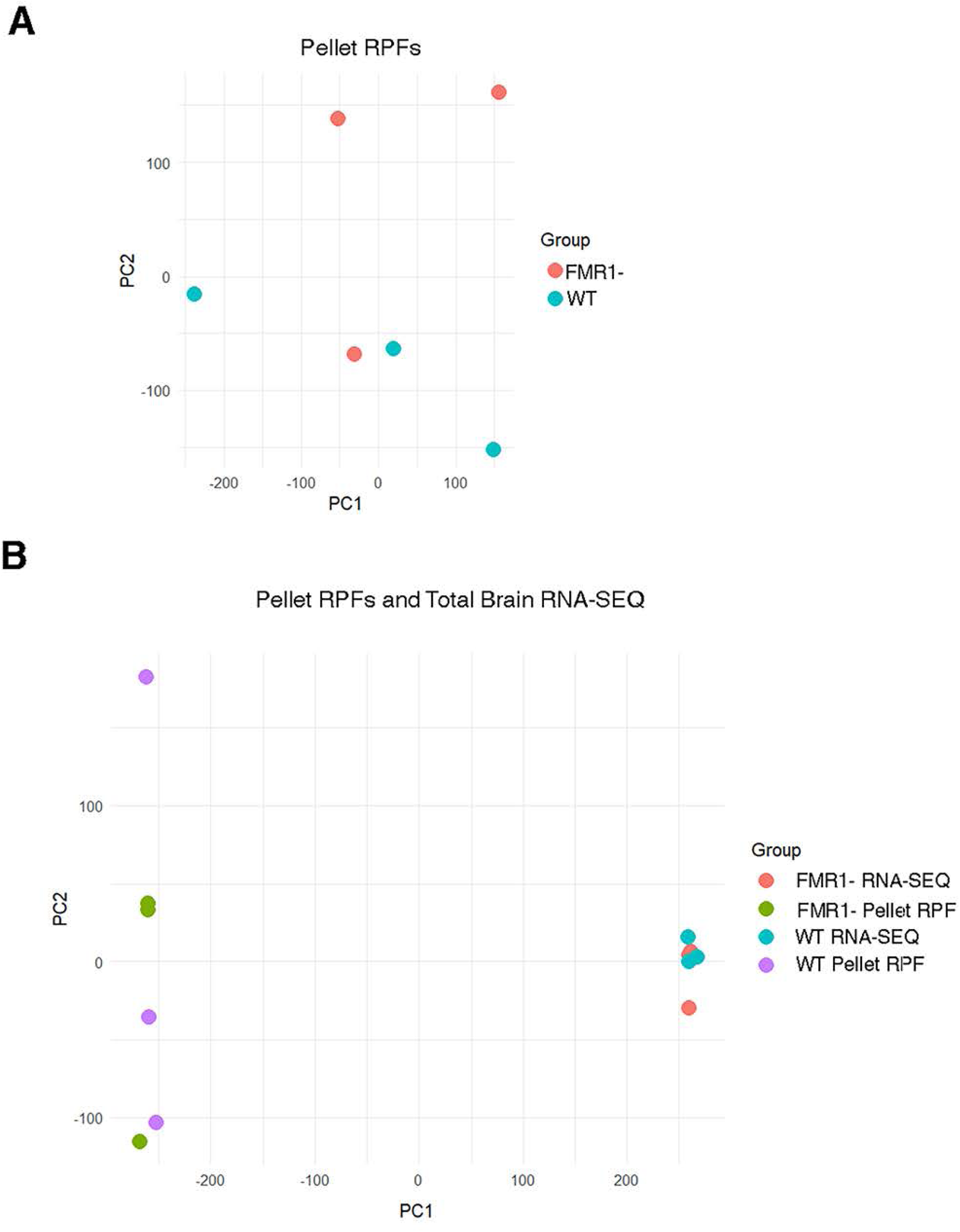
PCA Analysis. A) PCA analysis of Pellet RPFs from WT and FMR1-replicates. B) PCA analysis of Pellet RPFs and RNA-SEQ samples from WT and FMR1-replicates.

**Supplemental Figure S5-2:**
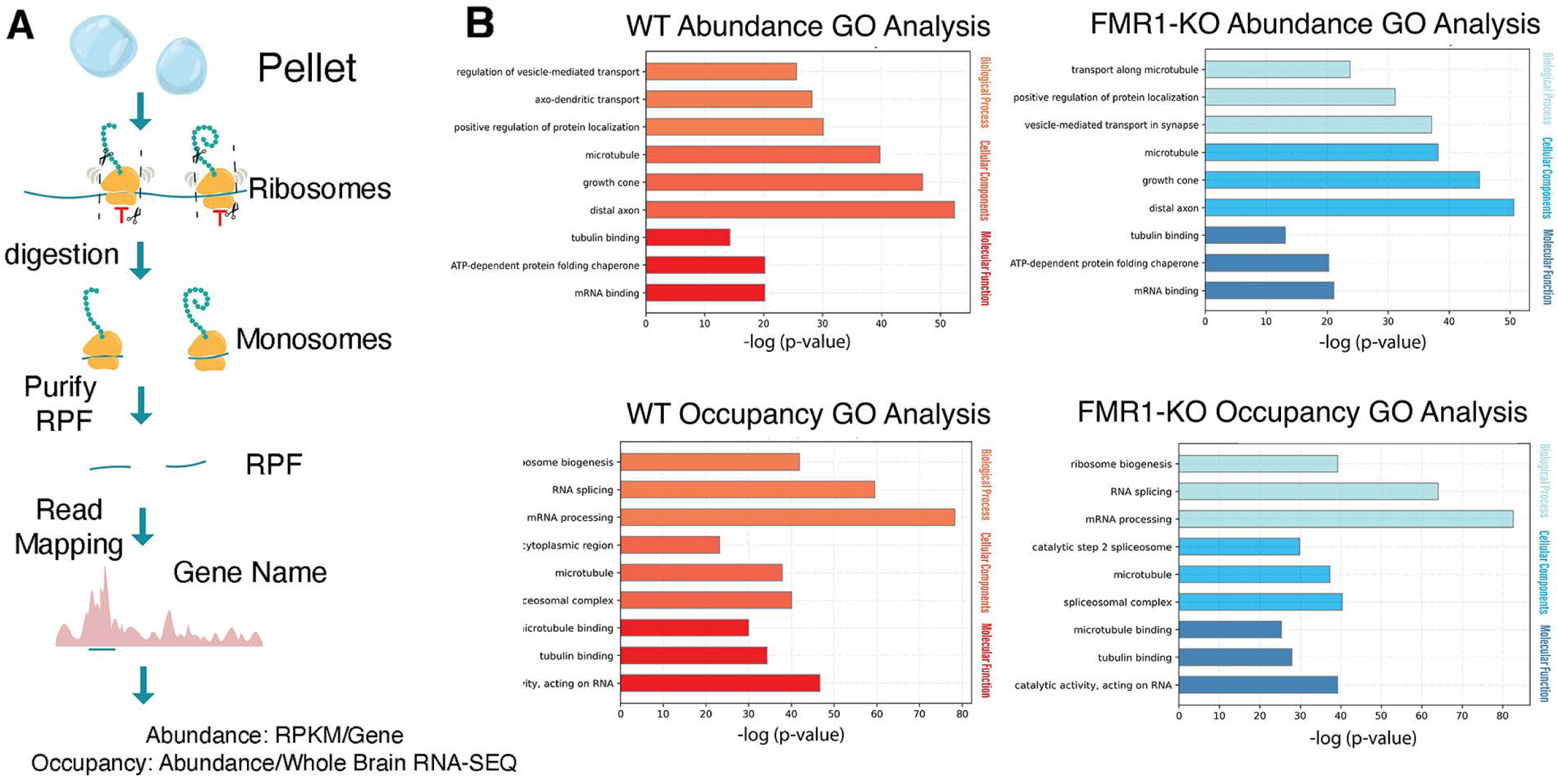
Assessment of RPF abundance, occupancy and enrichment in Pellets of WT and FMRI- mice. A) summary of protocol to generate RPF abundance, occupancy and enrichment. B) GO terms of the WT pellet (left) and FMR1- pellet (right)for abundance (top) and occupancy (bottom) For each graph, GO terms from the top 500 genes: Biological Function (top), Cellular Components (middle), and Molecular Function (bottom).

**Supplemental Figure 7-1.**
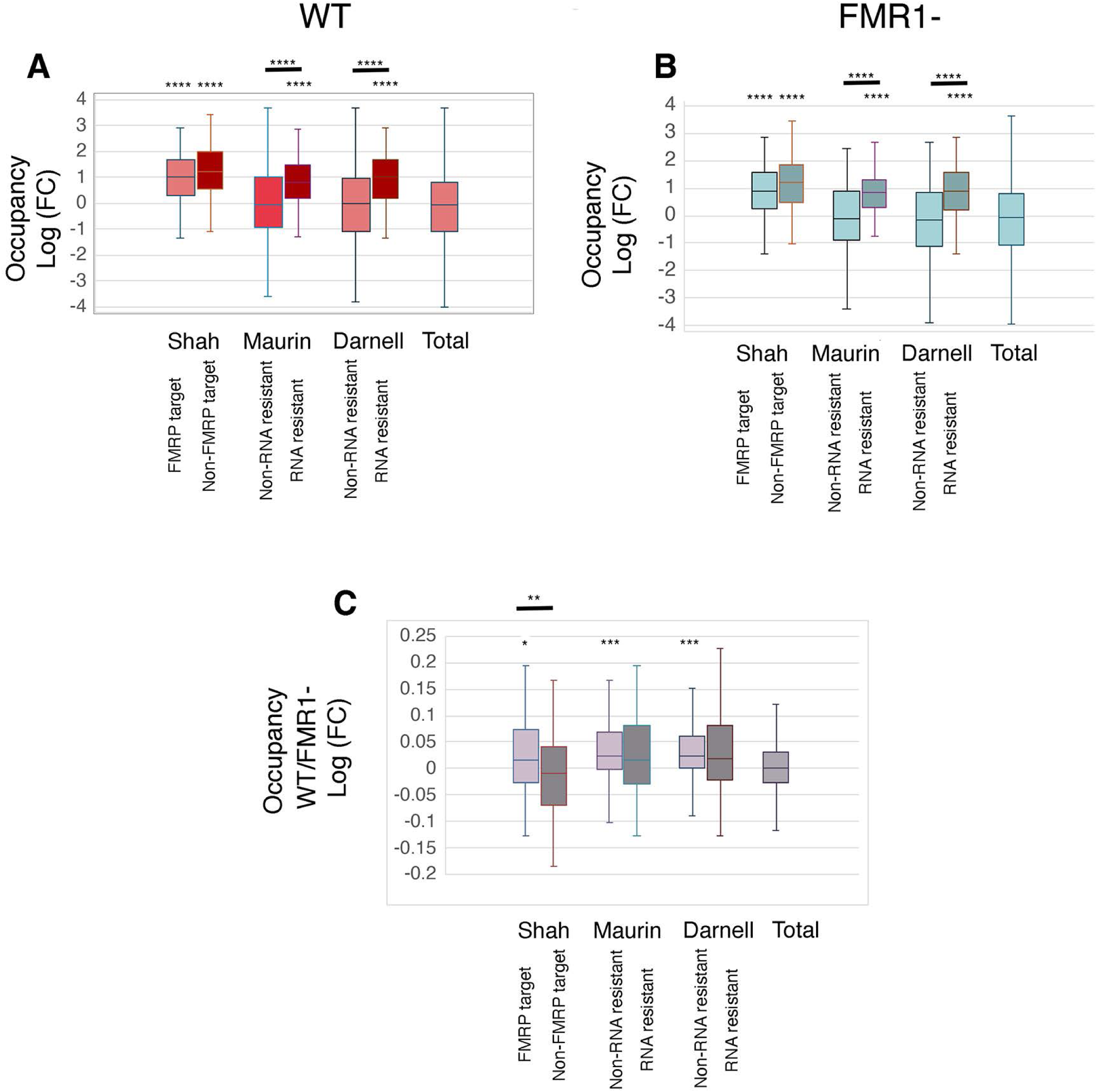
Occupancy analysis with Subsets of FMRP Targets.and RNAs resistant to Run-Off. The mRNAs found to be resistant to ribosome run-off (Shah et al, 2020) were divided into two sets consisting of mRNAs that are also FMRP targets (identified in Maurin et al, 2018 or Darnell et al;, 2011) or not. The FMRP targets were divided into two sets consisting of mRNAs identified as one of the 200 most abundant mRNAs resistant to run-ff (Shah et al, 2020) or not. Box and Whisker plots are shown with line for mean. Different from the total RNA group using two-tailed Welch’s t-test with Bonferroni correction for multiple T-tests. Differences between the two sets was also evaluated with a two-tailed Welch’s t-test. A) Occupancy in WT pellet B) Occupancy in FMR1- pellet C) Difference in Occupancy between WT and FMR1-

**Supplementary Figure S9-1.**
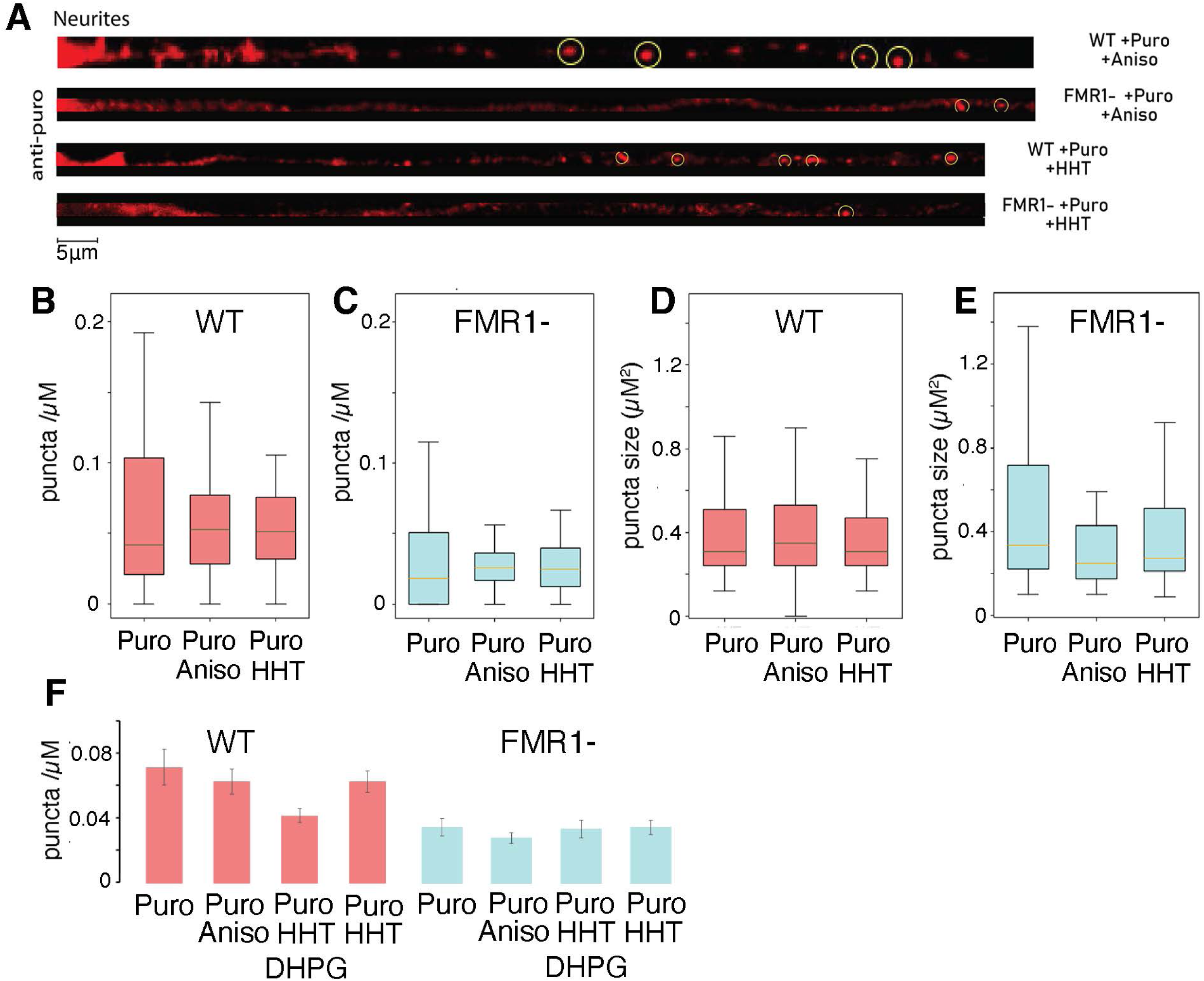
Effect of Anisomycin and HHT on RPM of hippocampal cultures derived from WT and FMR1-mice. A) Representative confocal images for puromycylated ribosomes treated either with puromycin and anisomycin together or pre-treated with HHT for 15 minutes before treating with puromycin. Circle denotes puromycin puncta. No visible staining was seen in the absence of puromycin. Quantification of RPM puncta density of puncta>50 microns from the cell body compared to WT (see Fig. 9) shown as box and whisker plits. There is no effect of adding anisomycin with puromycin or preincubation with HHT on the puncta density (B,D) or puncta size (C, E) in WT (B,C) or FMR1- (D,E). WT numbers are the same as Figure 9. Numbers are neurites/cultures. Puncta Density WT (42/5); WT anisomycin (A) (27/4); WT homoharringtonin (H) (36/5), FMR1- (41/4), FMR1-KO anisomycin (A) (23/3), FMRP homoharringtonin (45,5). One way ANOVA showed no significance for density or size (P>0.05). F) For clarity the data is also presented as mean +/- S.D.

## Supplementary Tables

**Supplementary Table S4-1.**
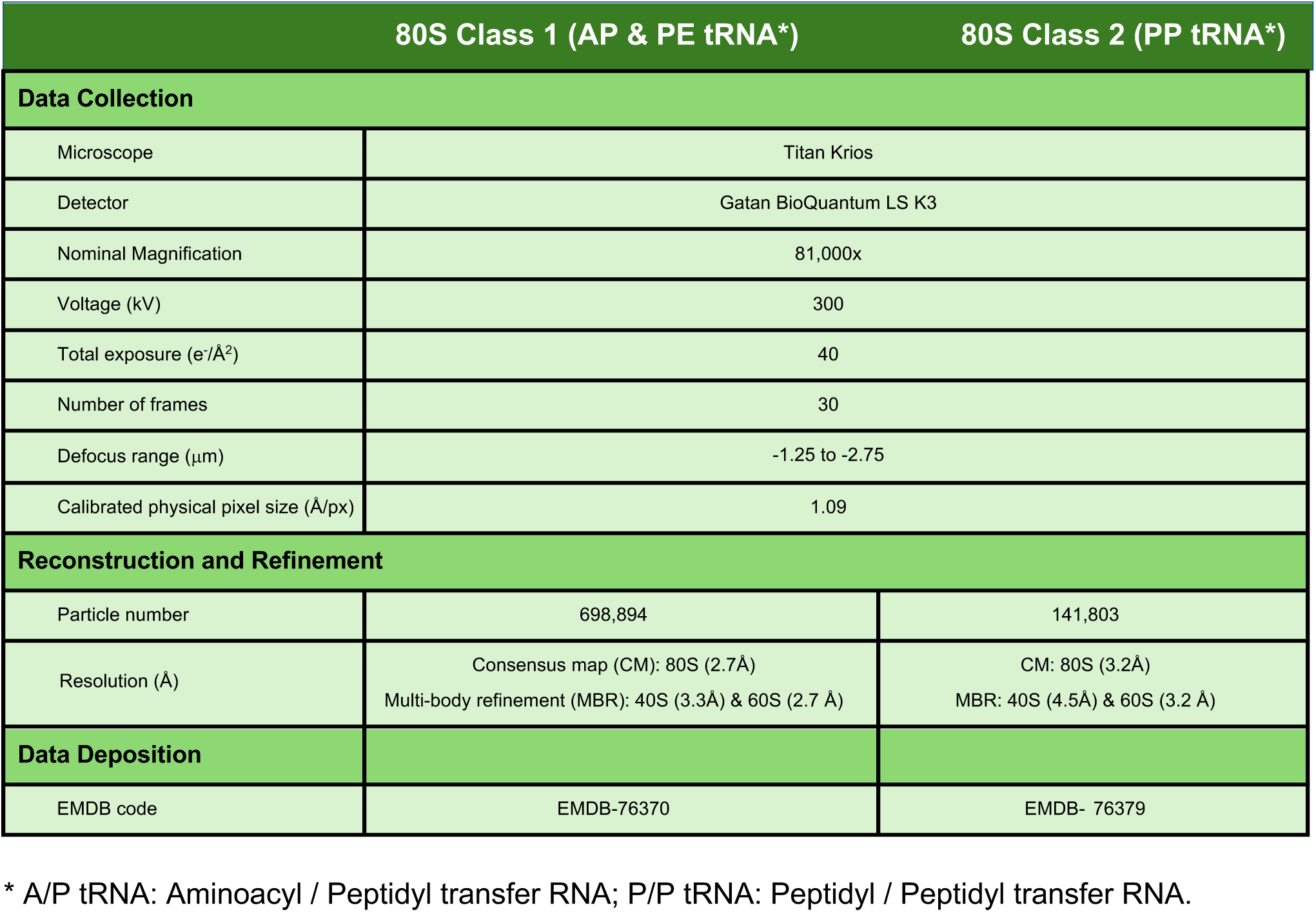
Data acquisition, reconstruction and refinement parameters and data deposition codes for the cryo-EM dataset.

